# Thermodynamics of cation binding to the sarco/endoplasmic reticulum calcium ATPase pump and impacts on enzyme function

**DOI:** 10.1101/507996

**Authors:** Bin Sun, Bradley D. Stewart, Amir N. Kucharski, Peter M. Kekenes-Huskey

## Abstract

Sarcoendoplasmic reticulum Ca^2+^-ATPase (SERCA) is a transmembrane pump that plays an important role in transporting calcium into the sarcoplasmic reticulum (SR). While calcium (Ca^2+^) binds SERCA with micro-molar affinity, magnesium (Mg^2+^) and potassium (K^+^) also compete with calcium (Ca^2+^) binding. However, the molecular bases for these competing ions’ influence on SERCA function and the selectivity of the pump for Ca^2+^ are not well-established. We therefore used *in silico* methods to resolve molecular determinants of cation binding in the canonical site I and II Ca^2+^ binding sites: 1) triplicate molecular dynamics (MD) simulations of Mg^2+^, Ca^2+^and K^+^-bound SERCA. 2) mean spherical approximation (MSA) theory to determine the affinity and selectivity of cation binding to the MD-resolved structures and 3) state models of SERCA turnover informed from MSA-derived affinity data. Our key findings are that a) coordination at sites I and II are optimized for Ca^2+^and to a lesser extent for Mg^2+^ and K^+^, as determined by MD-derived cation-amino acid oxygen and bound water configurations, b) the impaired coordination and high desolvation cost for Mg^2+^ precludes favorable Mg^2+^ binding relative to Ca^2+^, while K^+^ has limited capacity to bind site I, c) Mg^2+^ most likely acts as inhibitor and K^+^ as intermediate in SERCA’s reaction cycle, based on a best-fit state model of SERCA turnover. These findings provide a quantitative basis for SERCA function that leverages molecular-scale thermodynamic data and rationalize enzyme activity across broad ranges of K^+^, Ca^2+^ and Mg^2+^ concentrations.

## 1 INTRODUCTION

sarcoendoplasmic reticulum Ca^2+^-ATPase (SERCA) is a 110-kDa transmembrane cation pump which actively transports Ca^2+^ ions into the SR by utilizing energy released from adenosine triphosphate (ATP) hydrolysis [1]. SERCA has been widely studied for its role in returning intracellular Ca^2+^ to basal levels following stimuli that elevateCa^2+^ content [2]. The pump’s catalytic cycle is roughly characterized by four states comprising a sequential cycle *E*1 → *E*1*P*.2*Ca* → *E*2*P*.2*Ca* → *E*2. In E1, the Ca^2+^ binding sites are exposed to the cytosolic space, whereas the E2 conformations expose the low-affinity Ca^2+^ sites toward the SR lumen. The transition between E1 and E2 is driven by ATP hydrolysis at residue Asp 351 [1] following Ca^2+^ binding, for which E1P and E2P are the respective phosphorylated states of the enzyme. Accompanying transitions between catalytic states are prominent changes in its ten transmembrane (TM) helices as well as the cytosolic actuator (A) domain, nucleotide-binding domain (N) and phosphorylation domain (P). Many of these changes have been resolved through x-ray crystallography [3–6].

Based on available structural models of the protein and a wealth of biochemical studies [7–14], reactions schemes linking the E1 and E2 states are beginning to emerge. Utilizing SERCA vesicles coupled with spin label molecule, Inesi et al observed changed electron spin resonance spectrum upon Ca^2+^ binding that revealed conformational changes in the enzyme [11]. Dupont and coworkers similarly measured changes in intrinsic fluorescence upon Ca^2+^ binding and further proposed a two-step Ca^2+^ binding process to high affinity sites evidenced by the pumps slow rate of fluorescence changes [9, 10]. Additionally, conformational changes linking the E1 and E2 states were explored by kinetic studies of intrinsic fluorescence changes upon Ca^2+^ binding and release [7, 8]. To probe molecular determinants of Ca^2+^ binding in the pump, an E309Q mutant bound with two Ca^2+^ in its phosphorylated state was determined via x-ray crystallography [15], which revealed that the altered TM arrangements caused by the mutation leads to impaired pump functionality.With respect to cation binding affinities, Inesi et al measured Ca^2+^ binding and stoichiometry to SERCA vesicles in via chromatography [16], while others have probed the binding of the non cognate Mg^2+^ and K^+^ ions via intrinsic fluorescence changes in SERCA [17, 18].

**Figure 1:**
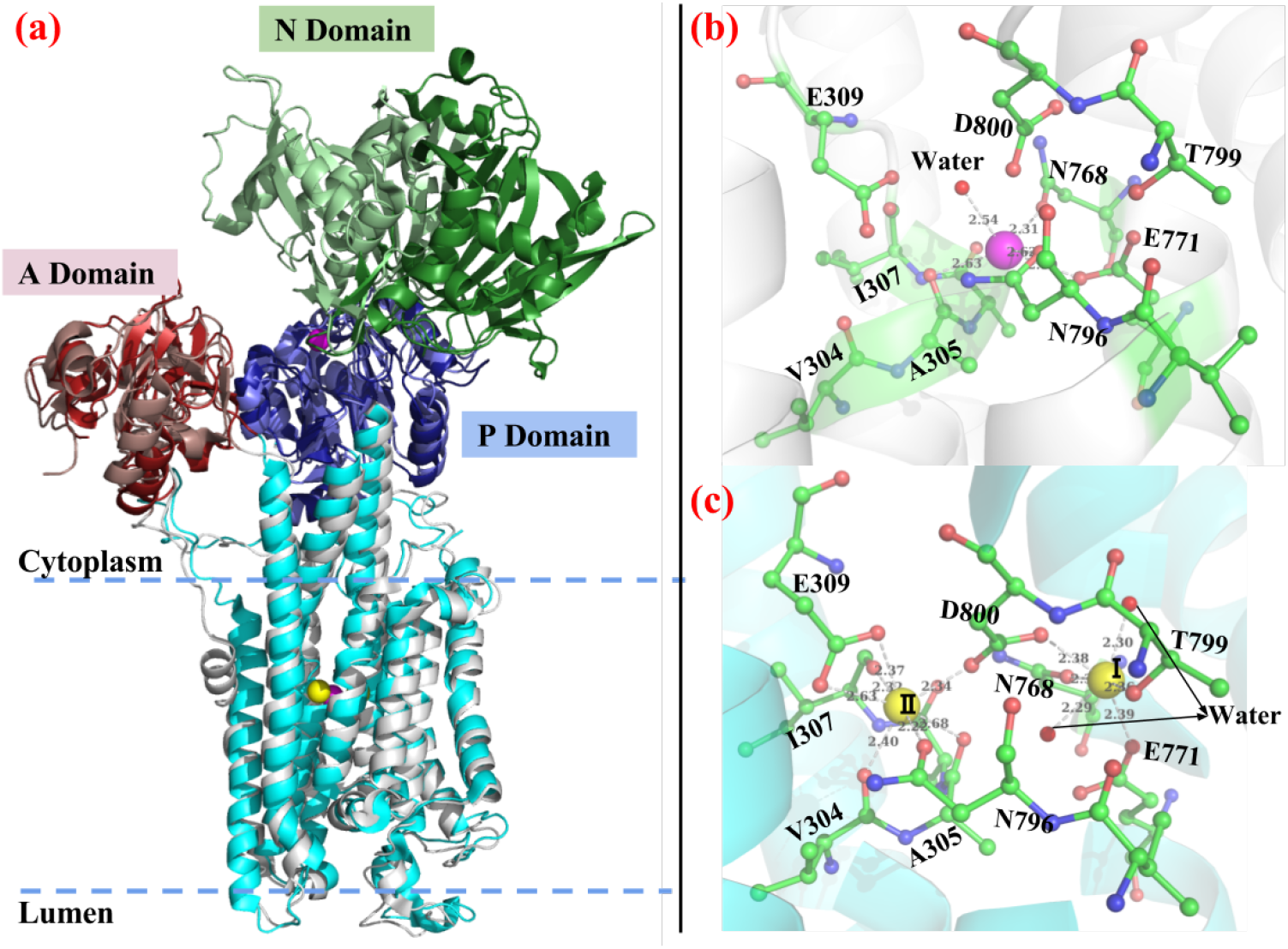
Comparison of Ca^2+^-bound (PDB ID: 1su4) and Mg^2+^-bound (PDB ID: 3w5b) SERCA crystal structures. (a) Superimpose of two crystal structures with 1su4/3w5b cytosolic domains (A, P and N) colored in red/salmon, blue/slate and green/palegreen. The TM helices are colored in cyan and grey for 1su4 and 3w5b, respectively. (b-c) Mg^2+^ and Ca^2+^ binding site comparison. In this orientation, sites I and II are right and left, respectively. Mg^2+^ and Ca^2+^ are represented as magenta and yellow balls. The distance between coordination oxygen atom and cation are also shown. The oxygen atoms of crystal water molecules are shown as red balls.

In complement to experiment, extensive molecular dynamics simulations have uncovered many aspects of cation binding and SERCA function that are difficult to probe via experimental approaches. Huang et al first explored the Ca^2+^ binding pathways to SERCA TM sites via MD simulation and observed the cooperative binding of two Ca^2+^ ions [19]. Kekenes-Huskey et al performed MD simulations on SERCA with and in the absence of Ca^2+^ to examine gating of Ca^2+^ binding by E309 [20], in addition to estimating ion binding free energies and kinetics. Espinoza-Fonseca et al evaluated the interaction energy between Ca^2+^/Mg^2+^/K^+^ and SERCA binding sites based on MD trajectories and reported that Ca^2+^ has the most negative interaction energy while K^+^ has the least negative value [21]. In addition to these initial studies on cation binding to the SERCA pump, more recent studies have probed mechanisms of SERCA function and its modulation by regulatory proteins and drugs [22–29].

Although studies have been reported on the roles of Mg^2+^, K^+^ on SERCA binding, less has been done to provide a thermodynamic basis for their effects on SERCA function. This is of particular importance, as intracellular K^+^ and Mg^2+^ concentrations are orders of magnitude larger than the roughly sub micromolar Ca^2+^ concentrations found in typical cells. Here, simulation studies of cation binding to molecular sites that resemble high affinity, amino acid-based motifs have been informative. Dudev et al for instance constructed cation binding sites using model compounds and calculated cation binding energies via quantum mechanics [30]. Ye et al designed a theoretical framework in combination with MD simulations to calculate cation binding free energies and isolate the energetic contribution from geometric and confinement effect [31]. Implicit models that describe Ca^2+^ binding thermodynamics as via ion density-based formalisms have also been reported, including reference interaction site modeling [32] and density functional theory [33]. Besides these methods relied on explicit binding site configurations, Nonner et al developed the MSA model in which the binding site is treated as confined filter filled with coordination oxygens. A variant of density functional theory called the mean spherical approximation (MSA) approach has proven effective in rank-ordering the binding of cationic species to oxygen-rich binding domains such as EF-hands in *β* parvalbumin (*β*-PV) [34] and Ca^2+^ channel selectivity filters [35, 36], through describing the electrostatics and hard-sphere contributions to the chemical potential of partitioning ions into oxygen-rich ‘filters’.

Our study has therefore focused on utilizing molecular dynamics (MD) derived data with thermodynamic and state-based models to assess contributions of Mg^2+^ and K^+^ binding on the SERCA turnover rate. Here we performed MD simulations of Ca^2+^, Mg^2+^ and K^+^-bound wild-type (WT) SERCA as well as the E309Q and N796A variants. These MD data provided structural information to assess cation binding thermodynamics via MSA to elucidate the molecular basis of SERCA’s preference of Ca^2+^ over Mg^2+^ and K^+^. Further, we relate these studies of E1 state ion binding to a state-based kinetic model of SERCA pump rate to determine the extents to which Mg^2+^ and K^+^ facilitate or inhibit catalysis. With this approach, we provide a multiscale and molecular basis for cation binding to SERCA and impacts on pump function.

## 2 RESULTS

### 2.1 Molecular dynamics simulations

We performed triplicate simulations of wild-type SERCA and its variants E309Q and N768A to probe the binding site coordination of the cations Ca^2+^, Mg^2+^ and K^+^, each replica was at least 100 ns in length. Ca^2+^- and Mg^2+^-bound configurations of the protein have been determined through x-ray crystallography, but to our knowledge, the binding of K^+^ to the Ca^2+^ binding domain has only been resolved via simulation [21]. In Fig. S1 we verify that the root mean squared deviations (RMSD)s of the peptide backbone atoms comprising the TM helices are relatively small (with Ca^2+^-bound cases around 1.1 Å and Mg^2+^/K^+^-bound cases around 1.5 Å) and do not significantly deviate from the input x-ray structures, suggesting that structures are near a local minimum. As shown in Fig. S2, the TM helices are quite similar across the cases considered, with minor deviations localized to the Ca^2+^-binding region. We note that the cytosolic A, N and P domains exhibit greater RMSD values (Fig. S3), which agree with previous analyses of the Ca^2+^ bound system [20, 21]. Although these domains were dynamic, they largely preserved the approximately 50 Å and 40 Å A-N domain distances in the input structures, as measured between the center of mass (COM) of the A domain (residues 1-47/123-240) and N domain (residues 358-603). Overall, the predicted Ca^2+^-bound variants and Mg^2+^-bound systems are consistent with previous structural [3, 37] and simulation [20, 21] studies.

We additionally report the root mean squared fluctuations (RMSF) of TM and cytosolic domain residues in Fig. 2 and Fig. 3, respectively as a qualitative gauge of the proteins’ stability. As shown in Fig. 2, the overall RMSF of all TM domains are small, with values of up to 3 Å attributed to TM3 in the WT Mg^2+-^bound case. In contrast, the Ca^2+^-bound cases yield the smallest RMSF values (approximately 1 Å), indicating that the TM regions are quite immobile when stabilized by bound Ca^2+^. However, the cytosolic domains yield larger RMSF values as shown in Fig. 3, most notably for the A and N domains (5 Å), which demonstrates significant mobility of cytosolic domains versus theTM region in agreement with previous studies [20, 21]. The domain motions are well-characterized as an opening/closing that we report as an A-N inter-domain distance (Fig. 3(d)). Our simulations generally reflected A-N distances that were close to the input structures in contrast to previous studies, which we attribute to the comparatively smaller simulation windows used in this study (100 ns versus > 500 ns).

**Figure 2:**
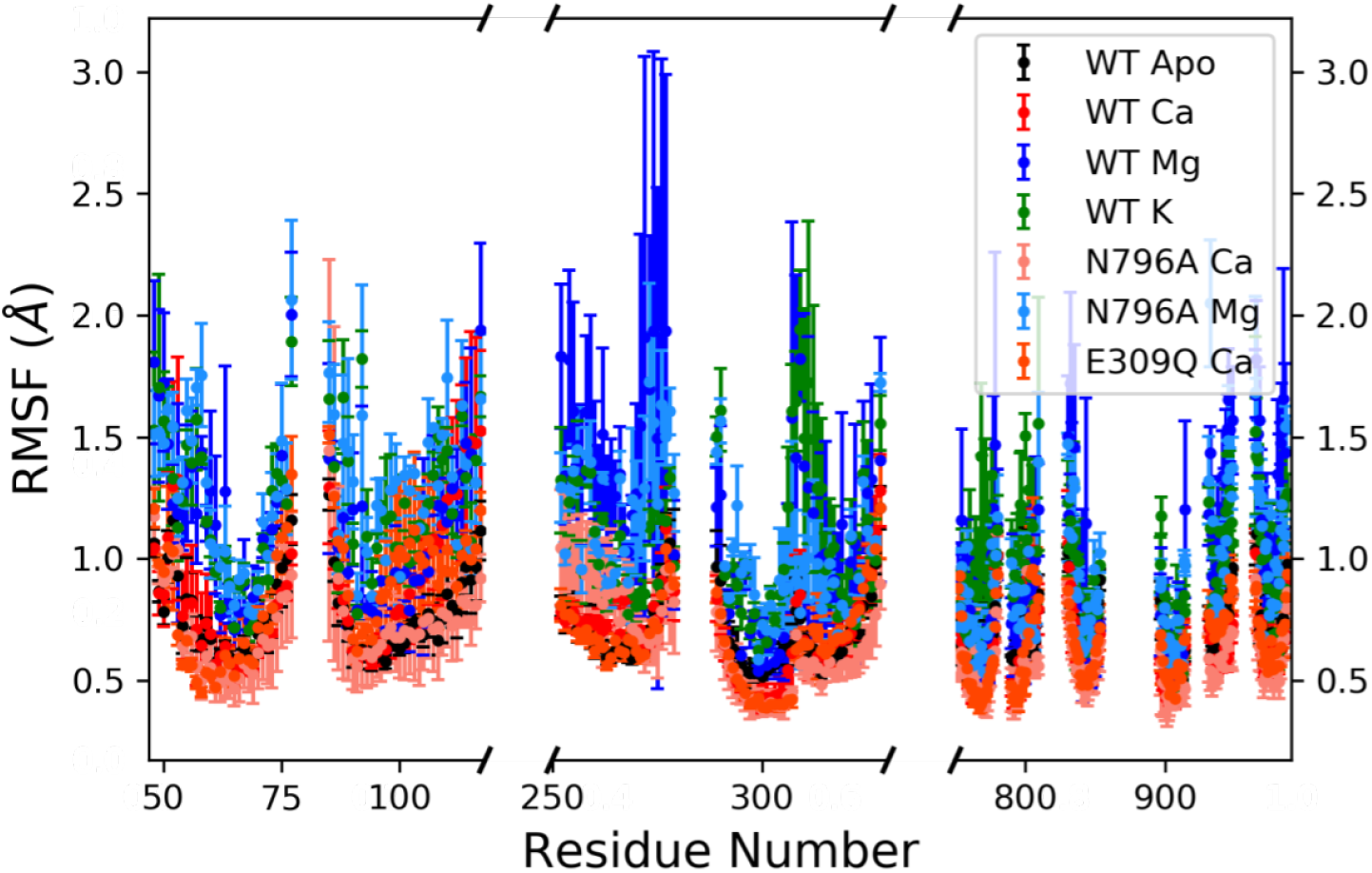
Heavy atom RMSF of each residue in the ten TM helices during the simulation. The error bars are calculated from the triplicate trajectories for each case. The gaps represent cytosolic domains which are plotted in Fig. 3.

**Figure 3:**
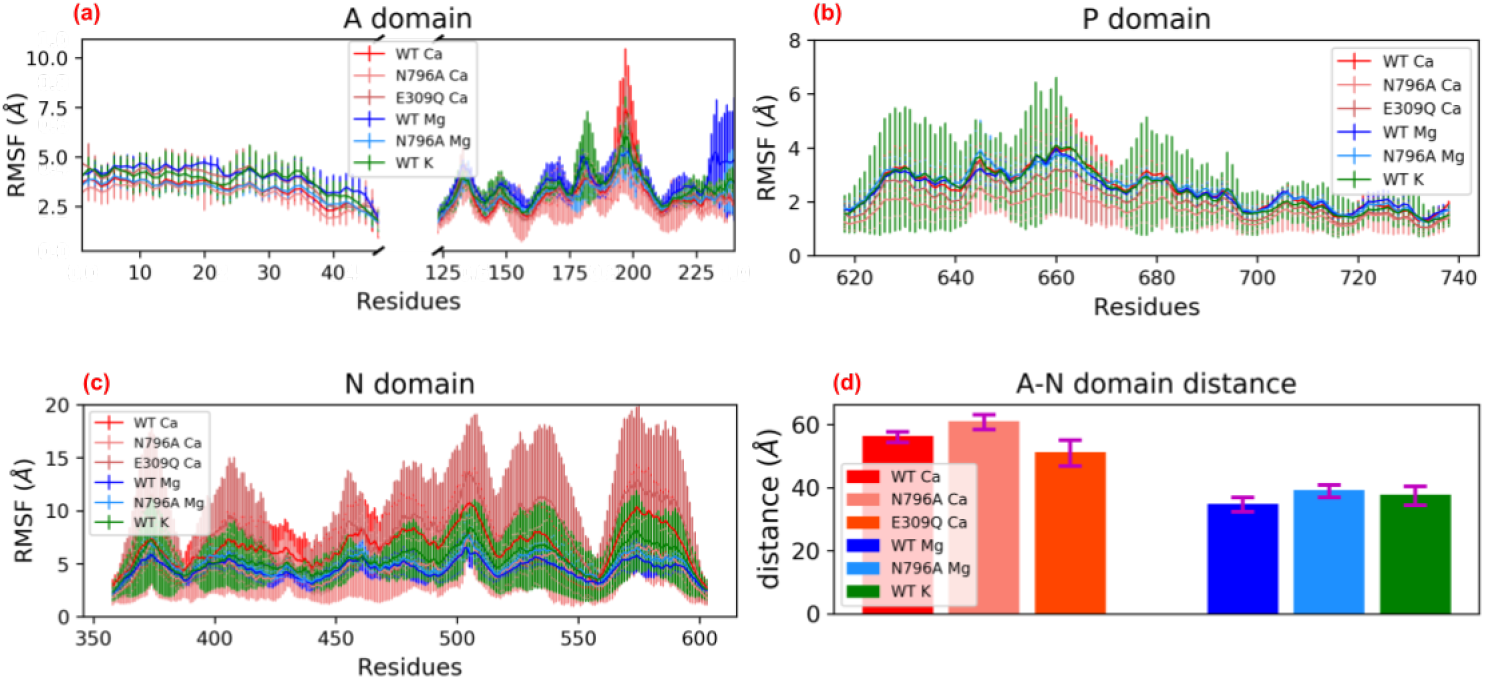
(a-c) Heavy atom RMSF of each residue in the cytosolic domain for WT and mutant SERCA during the simulation. The error bars are calculated from the triplicate trajectories for each case. (d) Distance between the COMs of A and N domains. The cytosolic domains exhibit greater mobilities compared to the TM helices.

For the K^+^-bound case, the simulations yield similar A-N distances as the Mg^2+^-bound cases, given that the Mg^2+^-bound x-ray structure PDB 3w5b served as the input structure for this configuration [21]. Overall, these RMSD and RMSF data are consistent with prior MD studies of WT SERCA [20, 21].

### 2.2 Cation coordination in the Ca^2+^ binding region

In this study, we highlight structural and dynamic contributions of the Ca^2+^ and Mg^2+^ binding domains to the thermodynamics of ion binding. This is analogous to our approach for probing Ca^2+^/Mg^2+^-binding to the *β* parvalbumin (*β*-PV) protein [34], for which we used MD simulation-derived structural data from the cation-bound configurations to parameterize a statistical mechanical model of fluid thermodynamics called mean spherical approximation (MSA). Specifically, we used the radius of the ions’ inner coordination sphere and amino acid oxygens comprising the sphere to estimate the binding site volume and coordinating oxygen density for MSA. We use a similar strategy for SERCA in that we assess the coordination of a given ion based on the number of oxygens within six Å of the bound ion. In contrast to our previous study, we additionally include coordinated waters in the MSA calculation that are directly involved in stabilizing the ion.

In Fig. 4 we report radial distribution function (RDF)s of cation-coordinating oxygens from the simulations. These RDFs demonstrate that Ca^2+^, Mg^2+^, and K^+^ have varying degrees of coordination with amino acid oxygens within the binding site. We summarize the identity of the coordinating amino acids in Sect. S1.4, although from the perspective of MSA theory, only the number of contributed amino acid oxygens is important. The Ca^2+^ within site I was optimally coordinated with 8.5 oxygens on average (including three water oxygens). Ca^2+^ within site II maintained six amino acid oxygens pairings, similar to the x-ray crystal structure, but did not directly coordinate waters. By virtue of having more coordinating oxygens, we speculate that the Ca^2+^ ion in site I is bound more tightly than that found within site II. In contrast, for the N796A mutant, the site I Ca^2+^ had a reduced coordination number of six relative to over eight in the WT structure. Interestingly, for the site II Ca^2+^ in the N796A variant, the loss of coordination to site 796 was compensated by interactions formed with E58 and a bound water molecule (see Fig. S11(b)) to yield a greater coordination number than observed in the WT. For the E309Q mutant, two possible side chain rotamers of Q309 were investigated, as both rotamers were viable starting positions (see Fig. S11(c-d) for illustration of rotatmer directions). We found that these rotamers yielded identical Ca^2+^ coordination patterns for the two binding sites: site I had six coordinating oxygens versus seven for site II, while neither included bound waters (see Fig. S11(c-d)). Similar to the N796A site II Ca^2+^ case, E58 in TM1 also participated into the coordination with Ca^2+^ at site II in both rotamers of E309Q. Overall, these simulations reveal that ion/oxygen pairing is remarkably labile between the sites I and II, and can incorporate bound waters to maximize Ca^2+^ coordination.

**Figure 4:**
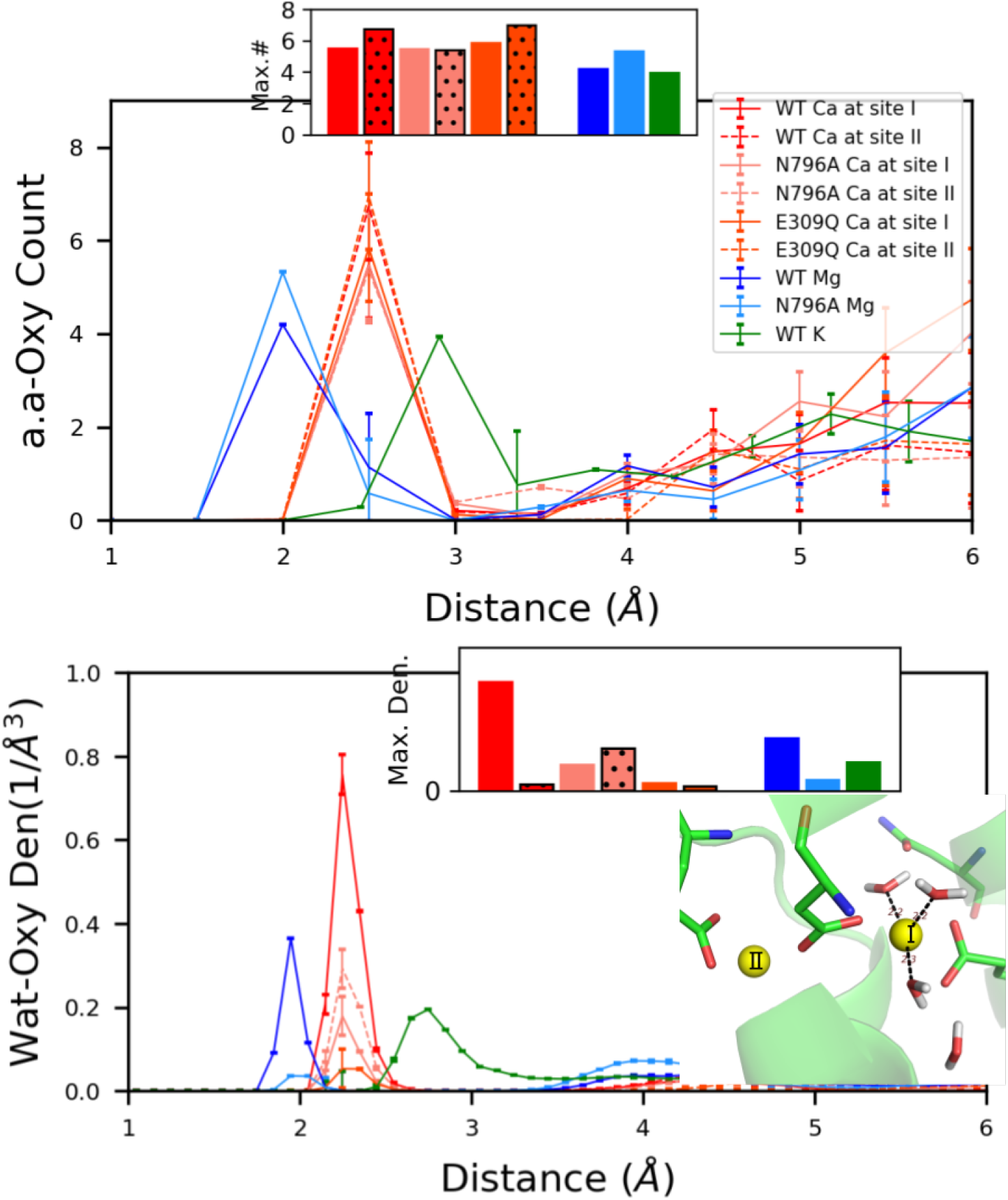
Radial distribution function (RDF) of amino acid and water oxygens atom about bound Ca^2+^, Mg^2+^, or K^+^. The distribution around Ca^2+^ for each individual case are shown in Fig. S9 for clarity. The insets bar graph show the maximum number of coordinating amino acid oxygen and water density around cation in each case (the bars with black dots represent site II Ca^2+^). The coordinating waters with site I Ca^2+^ of WT SERCA is also shown. Ca^2+^ ions bound to WT SERCA tend to reflect the highest degree of coordination among the modeled systems.

In contrast to Ca^2+^, Mg^2+^ binds in a ‘hybrid’ site between sites I and II with an average six coordinating oxygens (including one water molecule, see Fig. S13(a)). K^+^, on the other hand, binds site I with five coordinating oxygens (including one water molecule, see Fig. S13(c)), while K^+^ at site II is highly dynamic and interchanges with water immobilized in the SERCA interior. Given that these ions are both positively charged and not remarkably different in size relative to Ca^2+^, we had anticipated that the non-cognate ions might adopt higher coordination numbers in the native Ca^2+^ sites. Clearly the molecular simulations did not reflect this expectation; in Sect. 2.3 we provide a thermodynamic rationale via MSA theory for why these non-cognate ions present impaired coordination numbers.

The differences in cation-oxygen coordination patterns for the cases considered are accompanied by variations in the coordinating residues’ mobilities relative to WT. These mobilities are measured as RMSF values in Fig. 5, for which the upper row represents site I and the lower row, site II. Generally, Ca^2+^-bound systems have RMSF values for most residues around 0.5 Å, which are the smallest among the ions considered. To a certain extent, the reduced mobility could be interpreted as an indication of tighter and more favorable binding, although this would come at an entropic cost that is not explicitly estimated here. We were, however, surprised to see little change in RMSF for the Ca^2+^-free (apo) state versus the Ca^2+^-bound cases. One possible reason is that in apo state, waters fill the binding sites and stabilize residues via a hydrogen bonding network (see Fig. S7) - in this capacity, bound waters might ‘prop’ open the Ca^2+^-binding domains to promote rapid incorporation of solvated Ca^2+^ ions from the bulk medium. Additionally, we found that WT SERCA and its variants presented negligible differences in RMSF upon Ca^2+^ binding, whereas the non-cognate Mg^2+^ and K^+^ manifest significant RMSF increases across all residues (Mg^2+^ generally above 0.8 Å and K^+^ above 1 Å). It is possible that the greater mobility of coordinating residues for the non-cognate ions are indicative of impaired coordination. We had anticipated that waters could be incorporated into the ions’ binding domain to suppress fluctuations in amino acids contributing to coordination, much as was observed for the apo state. However, it is apparent the the strong electrostatic affinity for these ions with the coordination residues limited the volume within which waters could be incorporated. At a minimum, these data suggest that ion coordination is dynamic, with fluctuations on a nanosecond timescale (see Fig. S6), which ultimately may play a role in selecting Ca^2+^ over non-cognate ions.

**Figure 5:**
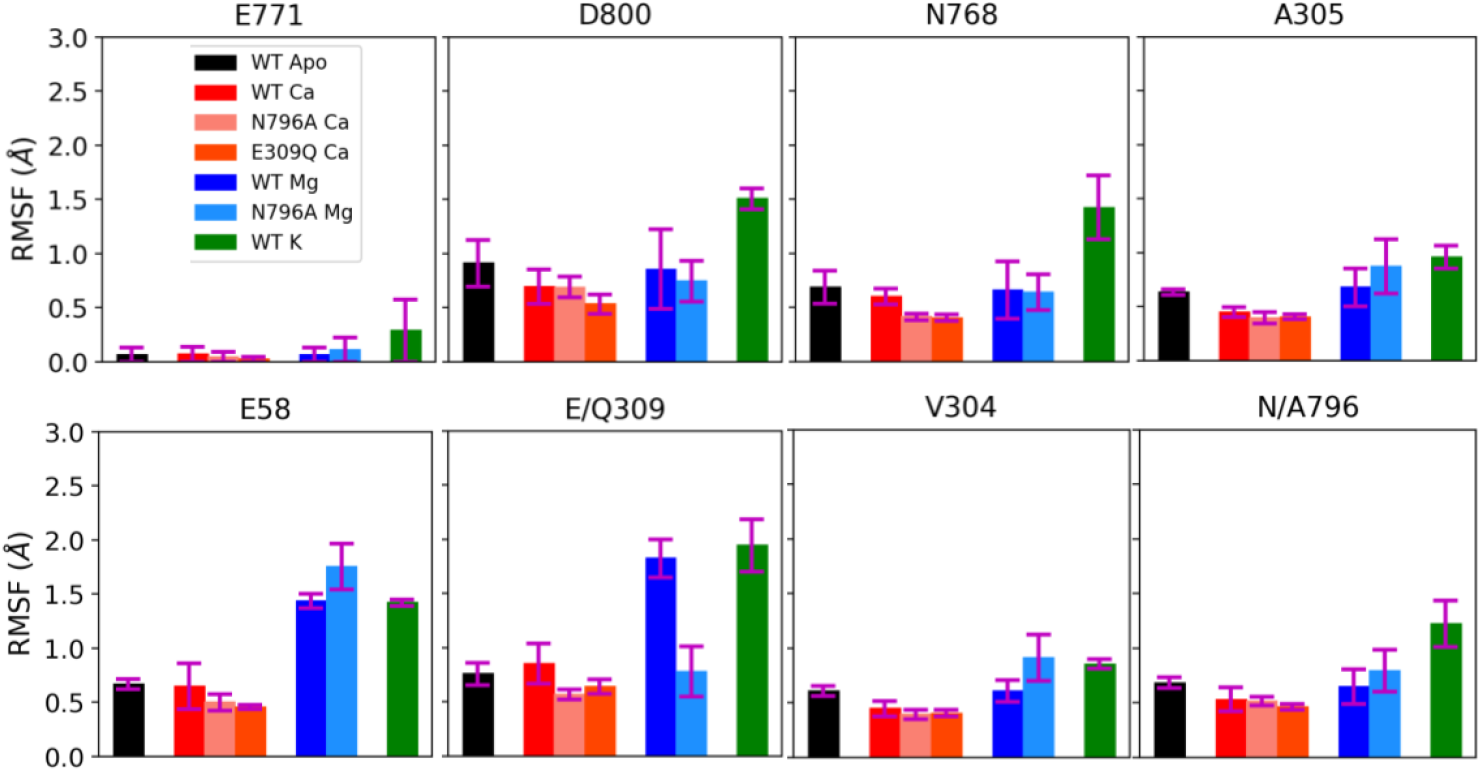
Heavy atom RMSF of key coordinating residues at the cation binding sites. The error bars were calculated from the triplicate trajectories for each case. The upper row represents site I residues while lower row were site II residues (E58 was not involved into coordination with cations in the crystal structure, although interactions with bound ions were predicted in our simulations, see Fig. S11). Ca^2+^-bound configurations generally exhibit lesser mobility compared to the configurations with non-cognate ions Mg^2+^ and K^+^.

### 2.3 Thermodynamics of ion binding at sites I and II

Molecular dynamics simulations provide qualitative insight to the binding of various ions in the binding sites of SERCA, but alone do not directly predict affinities. Therefore we explored MSA to semi-quantitatively estimate free energies and selectivity of ion binding. MSA predicts chemical potentials of partitioning solvated ions into the SERCA binding domains, assuming a similar approach that was performed for the *β*-PV [34] protein. Namely, MSA theory estimates the chemical potential of ion binding, based on the assumption of finite sized ions and chelating oxygens confined to a spherical binding site volume. To utilize this method, we first compute oxygen RDFs about bound ions. These data provide oxygen filter densities and volumes, from which chemical potentials of ion partitioning into the binding site ‘filter’ from the surrounding bulk solution can be estimated. As shown in Fig. 6, Ca^2+^ at site I of WT case presents the most negative and therefore thermodynamically-favorable MSA-predicted chemical potentials across all cases, corresponding to the largest number of coordination oxygens. For the SERCA variants, site I and site II Ca^2+^ ions have modestly less favourable chemical potentials compared with site I Ca^2+^ from WT cases, as the relative values were approximately 0.3 kcal/mol higher. These are consistent with the comparable Ca^2+^ coordination numbers present among variants. Compared with Ca^2+^-binding, Mg^2+^ binding at WT and mutant SERCA yielded more significantly disfavored chemical potentials at approximately 2 kcal/mol relative to WT site I Ca^2+^. Among all cases, the K^+^ relative potential was the most positive at 4.2 kcal/mol, indicating that K^+^ is the least thermodynamically favored at TM sites. For Mg^2+^ and K^+^, the MSA predicted potentials also correlated with the cation-coordination patterns as these two ions had reduced coordination number when compared with Ca^2+^ (5.2/4.7 for Mg^2+^/K^+^ versus 8.5 for site I Ca^2+^). These data show that MSA could capture the key factors governing cation affinities such as coordination number and binding site volume. In addition, with inclusion of coordinating waters in MSA, the predicted potential for WT site I Ca^2+^ is most favourable among all cases, which agrees with experimentally measured affinities. Our MSA results indicate that site I confers greater Ca^2+^ affinity due to extensive inclusion of water coordination, trends can be rationalized using MSA theory. In Sect. S1.4, we utilized GIST to assess the relative thermodynamics of water binding to the Ca^2+^ binding domain. In general, we found that when bound waters are present in the ions’ coordination shells, the predicted free energies are on the order of −12 kcal/mol and thus very thermodynamically favorable.

**Figure 6:**
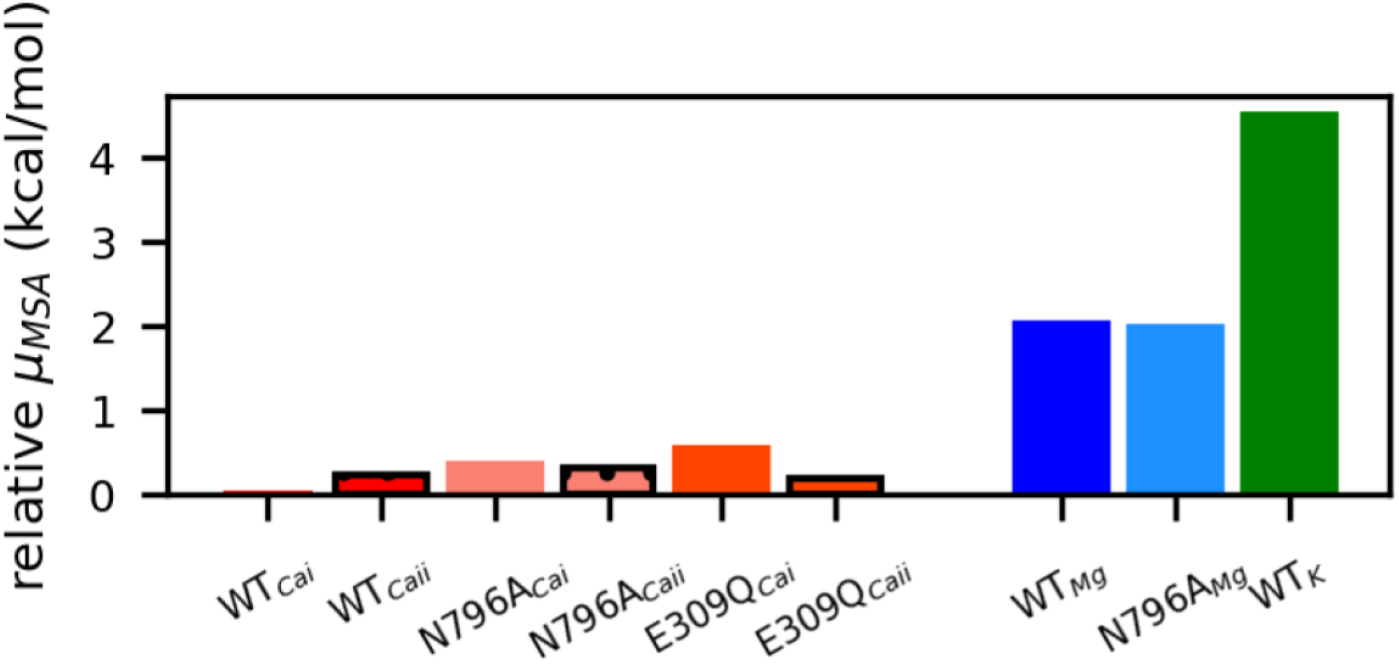
Mean spherical approximation (MSA)-predicted chemical potentials for each cation based on oxygen-coordination pattern (waters included) and optimal filter volumes derived from MD simulations. Potentials are given relative to Ca^2+^ bound to wild-type SERCA at site I. Ca^2+^ bound cases exhibit more favorable binding interactions compared to the binding of non-cognate ions, with the WT cases reflecting the most favorable potentials relative to the N796A and E309Q variants.

The membrane into which SERCA is embedded presents a negative surface charge owing to solvent-exposed phosphate head groups [38]. The corresponding negative electrical potential can attract positively charged ions and thereby increase their concentration near the transmembrane-bound SERCA, as we previously observed in [20]. Since the partitioning of cations into the SERCA binding sites depends on the composition of the surrounding electrolyte, we speculated that the local elevation of cations near the membrane surface would subsequently increase the concentration of bound cations. Additionally, the higher ionic strength could also reduce the desolvation energy and thereby further favor cation binding to SERCA, though this effect would likely be offset by screening electrostatic interactions between cations and the oxygens in the SERCA Ca^2+^ binding domains. To investigate this hypothesis, we determined the effective ion concentration near the membrane using a 1D solution of the linearized Poisson-Boltzmann equation [39], [*i*]_*eff*_ = *e*^−*βZiζ*^ [*i*]_*bath*_ where *Z*_*i*_ is the charge of ion and 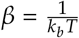 is ion concentration in the bath and *ζ* is the membrane potential.

Assuming *ζ* = −25 mV [40], we predicted that monovalent cation concentrations would be increased by 2.7 fold, anions would decrease by 0.37 fold, and the divalent Ca^2+^/Mg^2+^ ions would increase by 7.4 fold. As shown in Fig. 7, at low bath [Ca], both Mg^2+^ and K^+^ were present in the SERCA Ca^2+^ binding domain with concentrations of 7.7 M and 1.7 M, respectively. As bath [Ca] was increased, Ca^2+^ partitioned into the binding domain in favor of Mg^2+^; at roughly 2 × 10^−5^ M, the ratio of Mg^2+^ to Ca^2+^ was 1:1. The Ca^2+^ concentration at which Mg^2+^/Ca^2+^ was 1:1 varied by roughly 0.6 fold under the assumption of charged versus neutral membrane. In other words, according to our model, the local electrostatic environment about the membrane did not significantly impact the Ca^2+^ binding affinity. In contrast, in [20] we demonstrated that the negative surface charge densities of SERCA and the lipid enhanced the association rate of Ca^2+^ to the protein, thus the local electrostatic environment may have a greater contribution to ion binding kinetics than steady-state binding.

**Figure 7:**
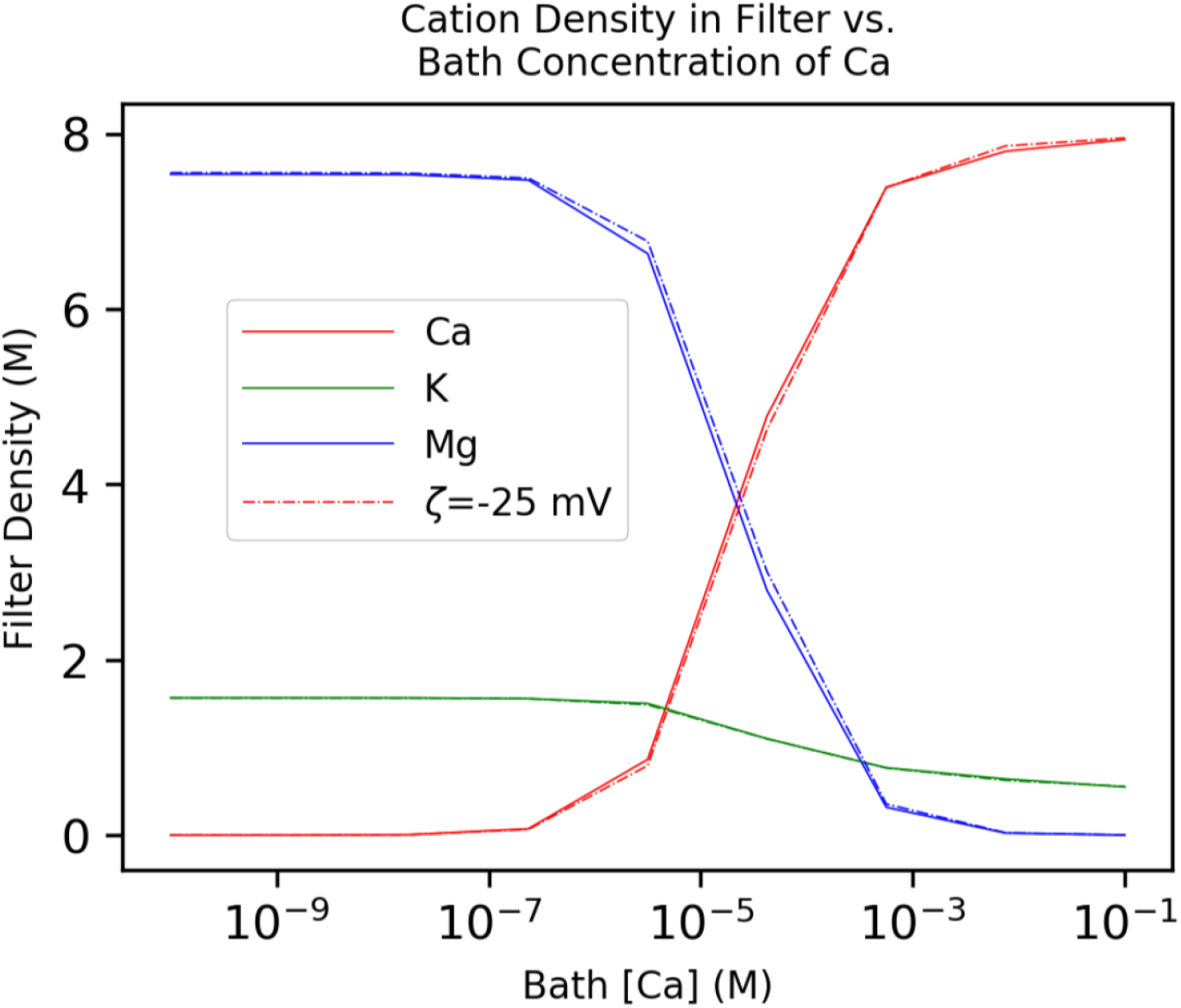
Cation concentration in the SERCA bind domains (assuming a filter volume = 0.33 nm^3^ from our MD simulations), *N*_*oxy*_ = 7 (coordination number of Ca^2+^ [30], this value lies between the coordination number of site I, *N*_*oxy*_ = 8.5, and site II, *N*_*oxy*_ = 6.7, for Ca^2+^ determined in our simulation of WT SERCA) and with the cation solvation energy from [33]. [KCl] = 150 mM and [MgCl_2_] = 2 mM. As cytosolic Ca^2+^ is increased, Ca^2+^ displaces Mg^2+^ bound to SERCA and reaches saturation at millimolar Ca^2+^ concentrations. Data are also presented assuming a membrane potential of *ζ* = −25*mV*, which locally increases bath cation concentrations by several fold according to Poisson-Boltzmann theory estimates.

### 2.4 Steady-state catalytic activity

Lastly, we relate our predictions of Ca^2+^ and non-cognate ion binding to the SERCA pumping rate. For the complete pumping cycle, two Ca^2+^ ions in the cytoplasm are transported into the SR by first binding SERCA to its E1 state. This binding process was proposed by Inesi et al [41] to consist of two Ca^2+^ successive binding events via a cooperative mechanism. Subsequent steps include binding of MgATP, a slow conformational transition to the E2 state, release of Ca^2+^ ions into the SR lumen, and a return to the E1 apo state. In practice, by accounting for the transition rates between SERCA conformational states, the time-evolution of each state can be described, which in turn can be related to the pump’s cycling rate. However, generally the transitions between ‘micro state’ conformations within the E1 or E2 ‘macro’ states, such as the Ca^2+^ binding steps in E1, are rapid relative to the slow E1 to E2 transitions. Hence, the micro states comprising the E1 and E2 stages are approximately in steady-state. This allowed us to describe the SERCA pump cycle rate as a two-state model for the E1 and E2 macro states (Eq. 8), which we used to relate experimentally and computation-determined binding constants to SERCA function.

In this state-based model, with exception to the undetermined transition rates between E1 and E2 (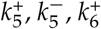 and 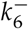), the resting rate constants and substrate concentrations were taken from [41, 42]. The above unknown rates were left as free parameters that were fitted to reproduce experimentally-determined turnover rates from [41] (see black data points in Fig. 8a)). This alignment was assessed as the difference between model predicted rates and experimental values reached minimum, as defined by

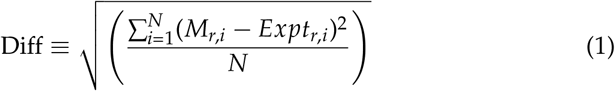

where *M*_*r*,*i*_ is state model predicted pump rate at experimental [Ca], *Expt*_*b*,*i*_ is the experimentally measured rate and *N* is the total number of data points. As shown in Fig. 8(a), the fitted Ca_*only*_ model (blue) reproduces the experimentally-determined SERCA turnover rates as a function of cytosolic Ca^2+^, which validates our state-based model.

**Figure 8:**
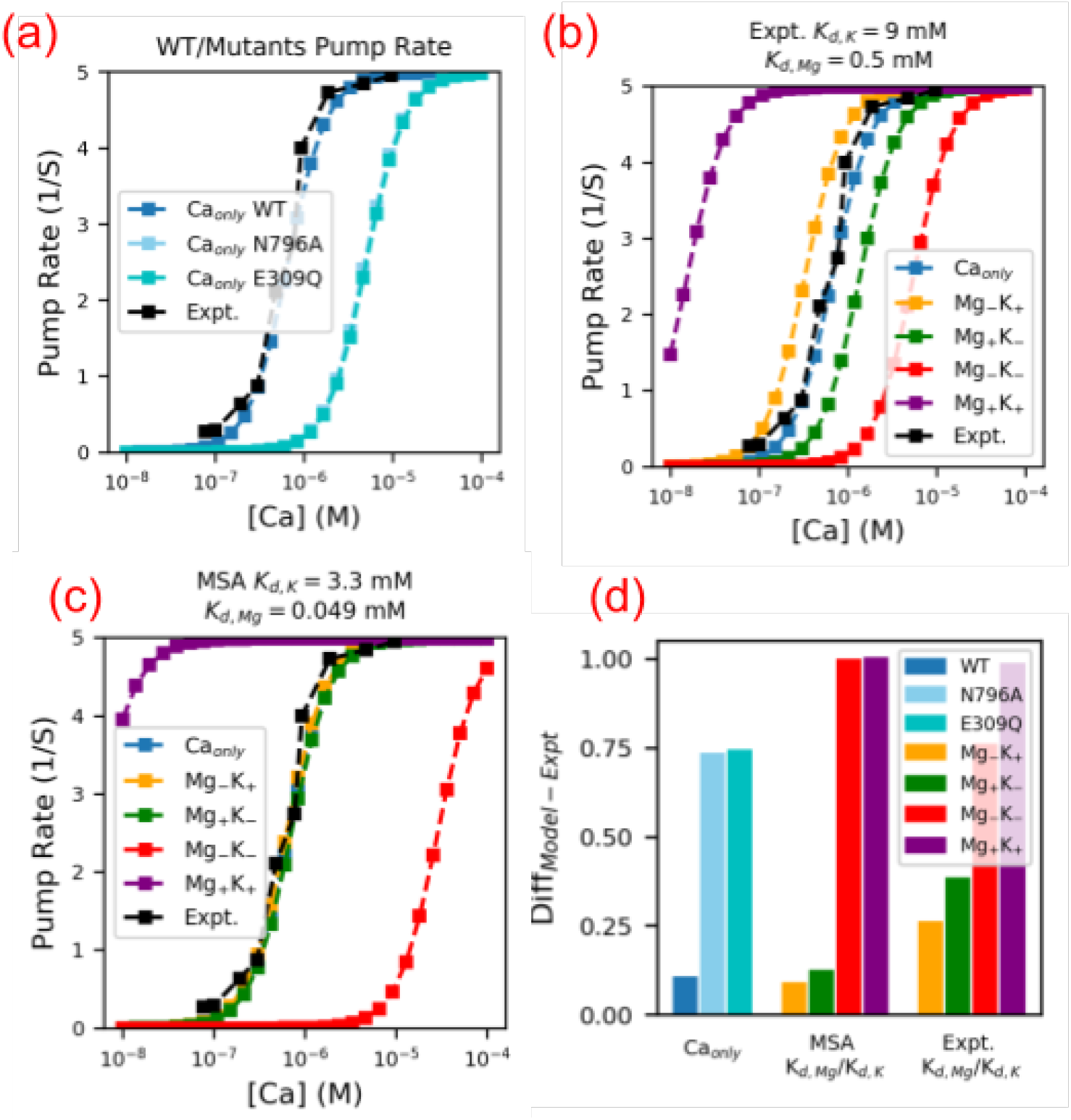
(a) Comparison of pump turnover rate between WT and SERCA variants using the Ca_*only*_ model with MSA predicted Ca^2+^ affinity for N796A and E309Q variants. The experimental data is from Inesi et al [41]. (b-c) Comparison of SERCA pump turnover rate of different state models against experimental data at [Mg] = 2 mM and [K] = 150 mM. In panel (b) the experimentally measured *K*_*d*,*K*_ and *K*_*d*,*Mg*_ in Table S4 were used. In panel (c) the MSA predicted *K*_*d*,*K*_ and *K*_*d*,*Mg*_ were used. (d) Normalized difference between each state model and experimental data for panels (a-c), as evaluated by Eq. 1. Our state-based models reproduce steady-state WT SERCA pumping rates and predict impaired rates for the N796A and E309Q variants.

### 2.5 Effects on non-cognate ions on steady-state behavior of SERCA

Since a primary focus of this study was to elucidate to which extent the noncognate ions Mg^2+^ and K^+^ influenced the SERCA transport cycle, we introduced additional microstates representing the Mg^2+^- and K^+^-bound configurations. The resulting representations are summarized in Fig. S4 and differ in terms of whether the ions serve as inhibitory or intermediates. Using the fitted model from Fig. 8a, we introduced dissociation constants for the Mg^2+^ and K^+^ species. We considered two strategies for defining those constants: 1) using experimentally-determined values from Table S4 and 2) constants determined from rescaling of the MSA-predicted values. For 2), the MSA predicted Mg^2+^/K^+^ chemical potentials were first converted to dissociation constants via 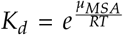. Second, the *K*_*d*_s were multiplied by a scaling factor, *λ*, that minimizes the difference between the MSA-predicted and experimentally-measured Ca^2+^ dissociation constants,

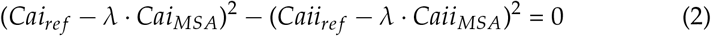

where *Cai*_*ref*_/*Caii*_*ref*_ and *Cai*_*MSA*_/*Caii*_*MSA*_ are site I/II Ca^2+^ dissociation constants from [41] (4 × 10^−8^ M and 4 × 10^−6^ M) and from MSA calculations (9.55 × 10^−6^ M and 1.38 × 10^−5^ M), respectively. Eq. 2 was minimized by *λ* = 0.17, thus yielding 4.93 × 10^−5^ M and 3.3 × 10^−3^ M for the MSA-predicted values of *K*_*d*,*Mg*_ and *K*_*d*,*K*_ for Mg^2+^ and K^+^ (see also Table S3). Predictions of the SERCA cycling rate using experimentally-determined dissociation constants are shown in Fig. 8(b) and the rescaled MSA constants in panel (c). Both approaches indicate that the Mg_+_K_−_ (green) and Mg_−_K_+_ (yellow) provide the best agreement with the experimentally-measured turnover rates, as these two models have relative smaller normalized difference values than other models, with the MSA-determined dissociation constants yielding the strongest agreement overall. Hence, the cycling rate data reported by Cantilina et al [41] was sufficient to eliminate two of the four proposed models.

To discriminate between the remaining Mg_+_K_−_ and Mg_−_K_+_ models, we next assessed the abilities of the respective models to reproduce steady-state Ca^2+^ binding data measured at various Mg^2+^ concentrations by Guillain et al [17]. In these experiments, both the E1.Mg and E1.2Ca states contributed a fluorescence signal indicative of Ca^2+^ saturation, therefore we report in Fig. 9 the combined probabilities of those states,

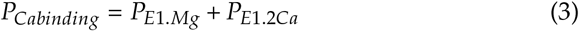

where *P*_*E*1.*Mg*_ and *P*_*E*1.2*Ca*_ are the probabilities of the E1.Mg and E1.2Ca states (see Sect. S1.3). The experimental data (black) shown in Fig. 9 demonstrate that Ca^2+^ saturation naturally increases with increasing cytosolic Ca^2+^, but importantly, saturation increases as Mg^2+^ is raised from 0 mM (circles) to 20 mM (solid triangles). These data indicate that Mg^2+^ locks SERCA into an E1 state in absence of Ca^2+^.We plot in Fig. 9(a-b) the predicted *P*_*Cabinding*_ data for the Mg_−_K_+_ and Mg_+_K_−_, respectively, as well as the fit in Fig. 9(c). We find that the Mg_−_K_+_ model provides the optimal fit with difference of 0.75 normalized to Mg_+_K_−_ model. The Mg_−_K_+_ model correctly captures the plateau in Ca^2+^ saturation at Ca^2+^ concentrations below 1 × 10^−7^, in contrast to the competing model. We note that as Mg^2+^ is increased to unphysiologically high concentrations (≥5 mM), the slope of the experimentally-determined saturation curves decreases, which is indicative of a loss in Ca^2+^-binding cooperativity. Our model does not directly consider ion-dependent modulation of Ca^2+^ binding cooperativity, therefore this behavior is not reproduced in our predicted data and accounts for some of the error relative to experiment. Additionally, we predict a greater population of the Ca^2+^-bound state at increasing Mg^2+^ levels than is experimentally observed, which accounts for the remainder of the error. Nevertheless, we find that Mg_−_K_+_ model provides the best agreement with experimental data, especially within physiological Mg^2+^ concentrations. Therefore, Mg^2+^ most likely acts as inhibitor and K^+^ as an intermediate in the SERCA pumping cycle.

**Figure 9:**
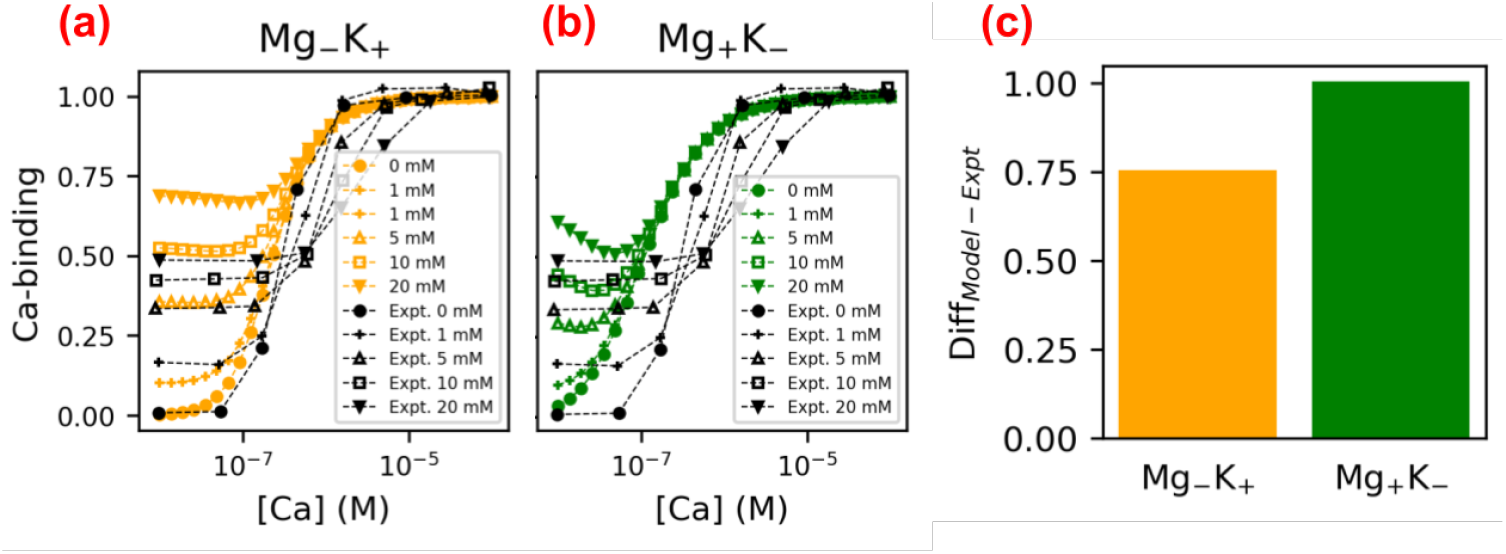
(a-b) Comparison of [Ca] dependence of Ca^2+^-SERCA binding of Mg_−_K_+_ and Mg_+_K_−_ models at varying Mg^2+^ concentrations ([K]= 150 mM) against experimental data from [17] Experimentally measured *K*_*d*,*K*_ and *K*_*d*,*Mg*_ were used. (e) Normalized difference between state models and experimental data as evaluated by Eq. 1. The Mg_−_K_+_ model, which assumes Mg^2+^ and K^+^ act as inhibitors and agonists, respectively, provide the best fit to experimental data.

Lastly, as a demonstration of how MSA-predicted Ca^2+^ affinities could be applied to infer changes in SERCA pumping function, we compared the turnover rates between WT and two variants using relative affinity estimates from Sect. 2.3. To be consistent with the procedure of obtaining Mg^2+^/K^+^ affinity from the MSA potentials, the same scaling factor *λ* = 0.17 was applied to the MSA-predicted affinities for the E309Q and N796A variants (Ca^2+^affinities at site I/II are 2.98 × 10^−6^ M/2.81 × 10^−6^ M for N796A and 4.06 × 10^−6^ M/2.25 × 10^−6^ M for E309Q). Relative to WT SERCA for which the half-maximal pump rate (approximately 2.5 per second) occurs at 6 × 10^−7^ M [Ca], the weaker affinities exhibited by the E309Q and N796A variants right-shift the half-maximal rate to ~ 1 × 10^−5^ M [Ca]. In other words, the SERCA variants are essentially nonfunctional within physiological Ca^2+^ concentrations (1 × 10^−7^ to 1 × 10^−6^ M [43]), which is consistent with experimentally-observed decreases in SERCA activity for the E309Q and N796A variants [44]. Although we recognize that the experimentally-measured activities arise from a culmination of factors beyond just the Ca^2+^ binding affinity in the binding domains, these data qualitatively indicate that MSA predicted affinities can then be used to rationalize steady-state pump turnover rates estimated from molecular-level simulations.

## 3 DISCUSSION

### 3.1 Ion coordination and contributions to cation binding affinity

A key contribution from this study was our use of MSA theory to evaluate trends in Ca^2+^ and non-cognate binding in the SERCA E1 states. By using molecular dynamics simulations of the wild type and two site-directed mutations, we could probe shifts in the binding site configurations - relative to the available crystal structures of the pump - that contribute to chelating cations. Overall, the MSA theory, when informed using molecular simulation data including water distributions, appears to be effective in rank-ordering ions by affinity (approximately −7 to −10 kcal/mol, see Fig. 10). Further, our state-based model of SERCA pumping function correctly captures cycling rates across physiological Ca^2+^ concentrations and predict functional effects of site-direction mutations (N796A and E309Q).

**Figure 10:**
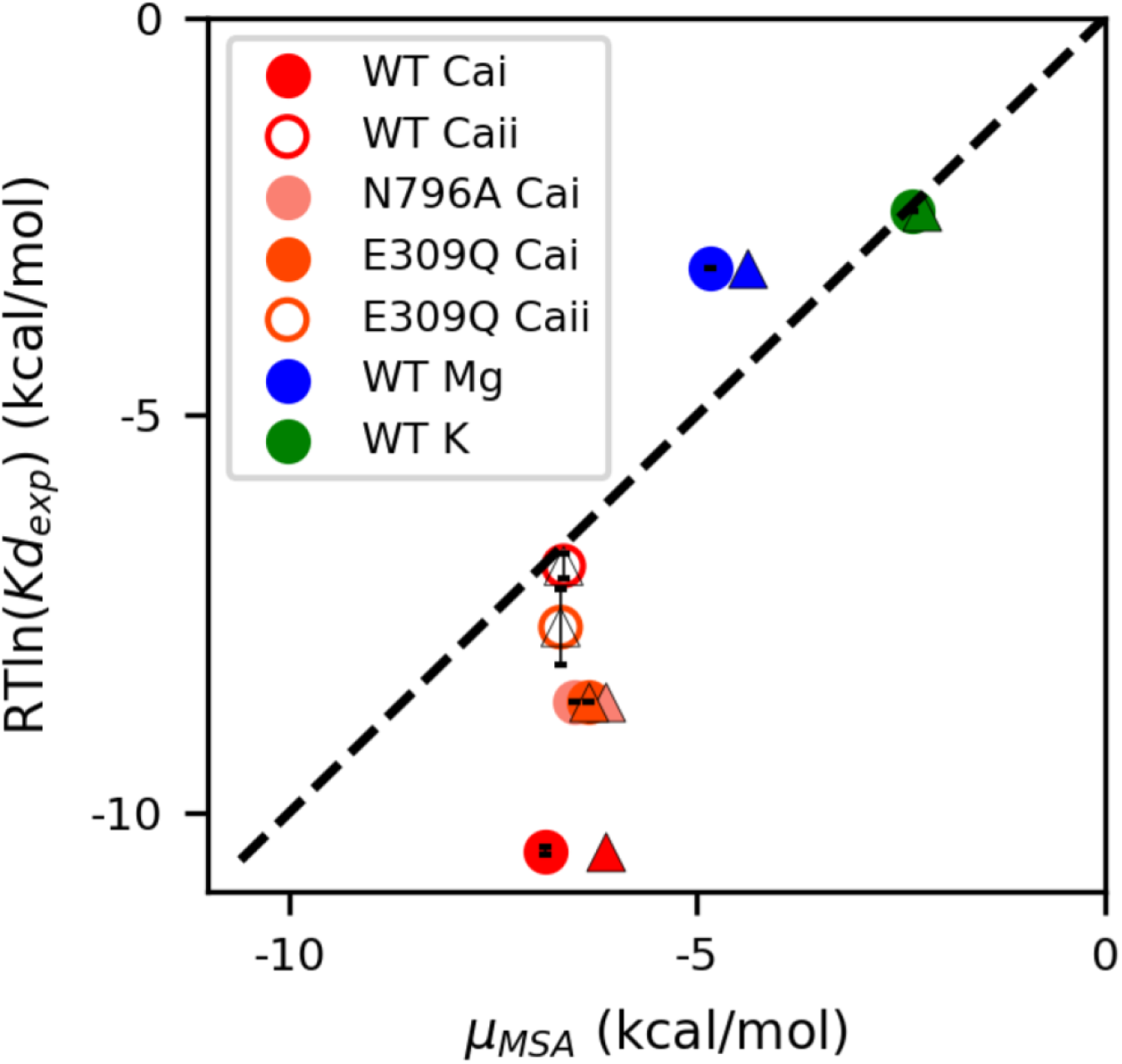
Correlation between MSA predicted chemical potential and experimentally measured binding free energy. The circles and triangles represent MSA results with waters and without waters taken into account, respectively. *RT* = 0.593 kcal/mol at room temperature and *Kd*_*exp*_ is experimentally measured dissociation constants in Table S4. The MSA chemical potentials correctly rank-order Ca^2+^ versus non-cognate ion binding.

We first applied this combined MSA/MD technique to an EF-hand containing protein called *β* parvalbumin to investigate factors contributing to its Ca^2+^ affinity and selectivity [34]. In both systems, the MSA/MD approach indicated that high affinity Ca^2+^ binding is afforded through the tight clustering of chelating oxygens with partial negative charges. Importantly, by predicting chemical potentials of ion binding, the MSA theory provided thermodynamic information about the relative competition between ions for the Ca^2+^ binding domains, which is generally unfeasible by MD alone. For instance, we determined that Mg^2+^, and not K^+^, is thermodynamically more probable to occupy the pump at low Ca^2+^ levels, but is disfavored as Ca^2+^ rises toward micro-molar concentrations typical in eukaryotic cells. In principle, this approach could additionally account for variations in electrolye composition expected near charged lipid bilayers, but we found scant differences in binding assuming a surface charge of 25 mV (see Fig. 7).

Consistent with our earlier findings for Ca^2+^-binding to *β* parvalbumin [34], MSA predicts increasing stability of the Ca^2+^ ion as the number of coordinating oxygens is increased. This trend reversed as the oxygen density increased beyond roughly six per 2.2 × 10^−1^ nm^3^, which is presumably when the volume is insufficient to accommodate all coordinating oxygens. For the E309Q and N796A variants, we potentially reduced the number of oxygens that could directly coordinate Ca^2+^. Based on the MD simulations for E309Q, however, it appeared that the loss of one carboxylic acid oxygen was compensated by a new interaction with E58 that maintained the same coordination number as the wild-type structure. This suggests that there is some degree of flexibility in coordination partners for the ion, which may facilitate the E309 residue’s role in kinetically gating the binding site [20]. In contrast, the N796A mutation was not offset by a nearby available oxygen, thus the predicted chemical potential was less favorable. **Therefore, our data suggest that the MSA could reasonably rank-order Ca**^2+^ **binding stabilities based on structures of the immediate coordination shell** based on Fig. 6, but is less sensitive to broad ranges of binding affinities reported in the literature (see Fig. 10).

We note that our predictions of favorable binding stabilities for Ca^2+^ at site II for the N796A and E309Q variants are at odds with Inesi et al [45, 46], who suggested based on SERCA ATPase activity that the site was incapable of binding Ca^2+^. Surprisingly, a structure of the E309Q variant (PDB ID: 4nab [15]) deposited in the protein databank contains Ca^2+^ at both binding sites. Ostensibly, the E309Q variant has a much lower, but nonzero affinity for Ca^2+^. Thus, it is likely that the MSA model does not sufficiently penalize nonoptimal Ca^2+^ coordination, or reflect changes in internal strain that might disfavor coordination [34].

We also found that including water distributions made a modest improvement in chemical potentials, as per Fig. 10. Based on Fig. S5, the narrow binding site volumes presented in sites I and II favor Ca^2+^ and Mg^2+^ over K^+^, while both divalent ions are increasingly stabilized by greater densities of coordinating oxygens and waters. It is interesting that the water-free MSA calculations indicate Ca^2+^ could be more stable in a binding site volume of 2.2 × 10^−1^ nm^3^, which is smaller than the volume presented in the MD simulations, albeit with a fewer number of oxygens (approximately five versus six). However, when water is considered, the stabilities at the smaller binding volumes are disfavored. It is possible that there is a kinetic advantage to having mobile waters involved in coordination, which could favor more rapid exchange of bound ions with the surrounding solvent. It is also possible that the trend can be explained on a thermodynamic basis, namely that by preserving waters that coordinate in the ion in bulk solvent, the desolvation cost upon binding SERCA are reduced, which should increase the apparent binding affinity. An intriguing possibility is that the Ca^2+^ affinity could be indirectly tuned by controlling the density of binding site waters.

The primary function of the SERCA pump is to transport cytosolic Ca^2+^ into the sarcoplasmic or endoplasmic reticulum, therefore we verified that MSA would indeed predict an unfavorable change in chemical potential based on conformational changes induced in sites I and II upon forming the E2 state. Since the focus of this study was on the E1 Ca^2+^ binding thermodynamics, we did not perform MD simulations of the E2 state. Nevertheless, based on the crystal structure of SERCA/beryllium fluoride complex(PDB ID: 3b9b [6]), which represents the pump’s E2P state, we illustrate in Fig. 11 that drastic changes in the Ca^2+^ binding site configuration culminate in a significant reduction in oxygens that could potentially coordinate Ca^2+^. We further evaluate the binding stability for a hypothetical Ca^2+^ bound between the labeled coordination groups and find that the MSA values are not only more positive than those of the E1 state, but are additionally greater than 0 kcal/mol. The positive values of approximately 1 kcal/mol indicate that Ca^2+^ binding at these position is less thermodynamically favorable than partitioning into the surrounding bulk electrolyte. In other words, when SERCA transitions into the E2 state, it is thermodynamically preferred for Ca^2+^ to vacate the binding site in favor of the reticulum lumen. Along these lines, mutations that alter the free energy difference for the Ca^2+^ sites in the E1 and E2 configurations could affect the efficiency of the ATPase.

**Figure 11:**
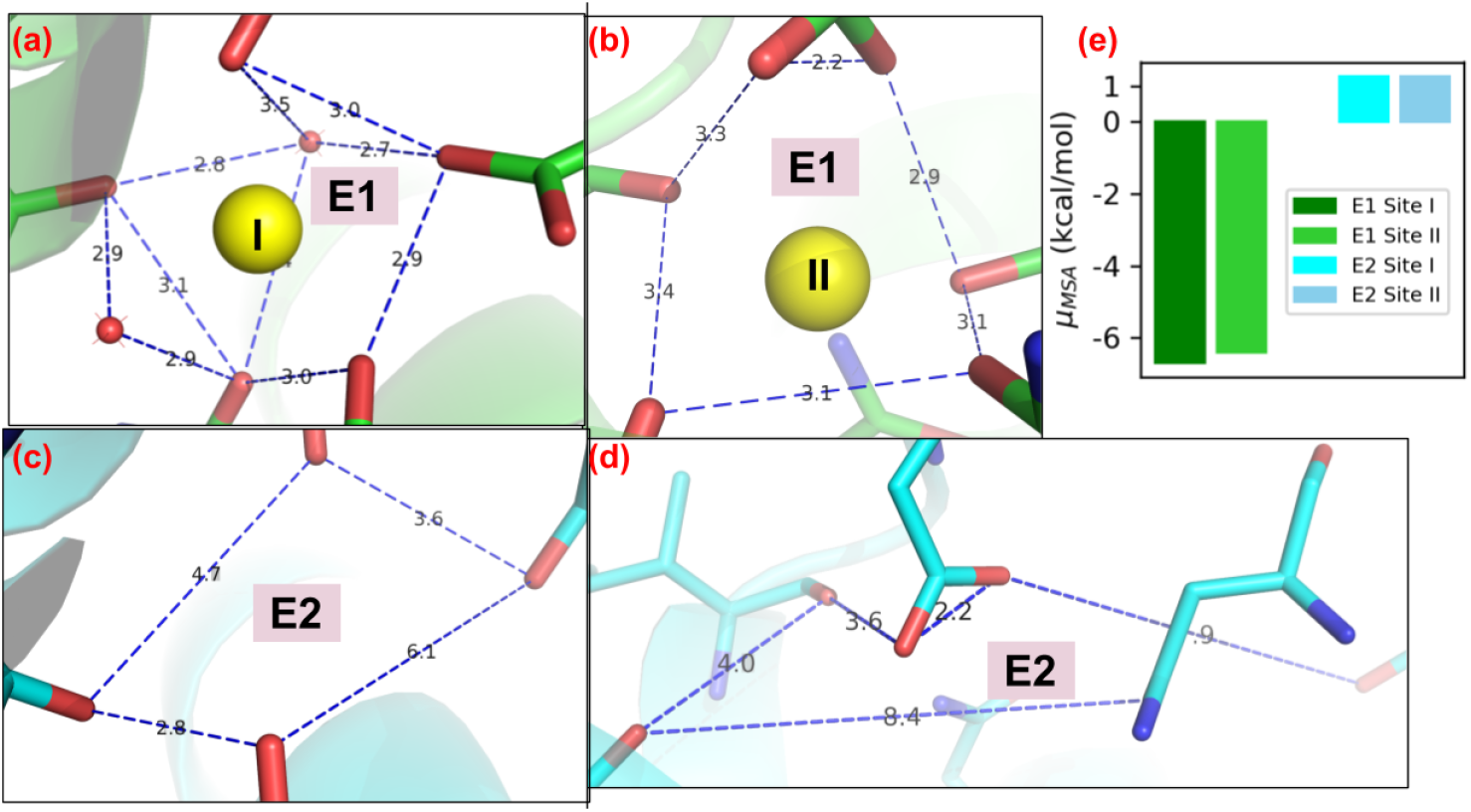
Comparison of Ca^2+^ binding sites in E1 state (a-b) and E2 state (c-d) from crystal structures of PDB 1su4 and 3b9b [6], respectively. Ca^2+^ and water molecules are shown as yellow and red spheres, respectively. The blue dashed lines in (a-b) outline the overall shape of Ca^2+^ coordinating spheres in E1 state while in (c-d) they depict assumed Ca^2+^ binding sites in E2 state. Panel (e) depicts MSA predicted Ca^2+^ chemical potential for these two sites. In E1 state the number of coordinating oxygens are seven and six for site I and II while these values are assumed as four for both sites in E2 state.

It has been suggested that the number of cations (Ca^2+^, Mg^2+^, and K^+^) bound to SERCA is approximately constant across its many conformational states [47]. In other words, Ca^2+^ binding is offset by Mg^2+^ and K^+^ dissociation. ATP is bound to SERCA in complex with Mg^2+^ [48], but there also exists structural [37] and activity [49] data that confirm Mg^2+^ binds in the pump’s transmembrane region. Toyoshima et al [37], for example, obtained the x-ray crystal structure of the pump with a single Mg^2+^ bound at a ‘hybrid’ trans-membrane site, for which the ion is coordinated by ~5 oxygens with distances of approximately 2.0 Å. Since Mg^2+^ bears the same charge as Ca^2+^, but with a modestly smaller radius, it is somewhat surprising that Mg^2+^ preferrentially binds at an intermediate site between the canonical Ca^2+^ binding sites I and II. We attribute the thermodynamic preference for Ca^2+^ at those sites based on two factors: the higher desolvation energy for Mg^2+^ relative to Ca^2+^ (779.94 kT vs 648.65 kT [50]) and the smaller binding site volume for the intermediate site versus sites I and II. With respect to the latter factor, in principle, Mg^2+^ could reduce the site I and II volumes by pulling the chelating oxygens toward the smaller ion, as we previously observed in [34]. For the EF-hand (helix-loop-helix) containing protein, we found that constricting the binding loop region introduced greater strain for Mg^2+^ relative to Ca^2+^, which we suggested would reduce the overall binding affinity for Mg^2+^. Although we did not explicitly evaluate strain that could be introduced upon Mg^2+^-binding for SERCA, we anticipate an even greater cost for reducing the binding site volume, since the chelating amino acid are firmly tethered to relatively immobile transmembrane helices. It is important to emphasize, though, that the non-optimal volumes for Mg^2+^ binding at sites I and II does not preclude the ion from binding, as we demonstrate high binding probabilities at very low Ca^2+^ levels (see Fig. 9).

Consistent with our earlier findings in *β*-PV, K^+^ is disfavored from binding the native Ca^2+^ binding volume based on its significantly larger radius and smaller charge [51]. However, K^+^ and other monovalent cations have been demonstrated to stimulate SERCA function, with K^+^ being the most efficient agonist [18, 47, 49]. Moreover, based on molecular simulations performed in the presence of transmembrane-bound K^+^, Espinosa-Fonseca et al suggest that K^+^ facilitates the pump’s E2 to E1 transition through its stabilization of the E1 state [21] In our simulations, we found that of the two K^+^ ions placed in the Ca^2+^ binding domains, the site I K^+^ remained bound with approximately four coordinating oxygens with ion/oxygen distances exceeding 4 Å. Based on the MSA predictions, although K^+^ has a favorable (*μ* < 0) binding potential that is consistent with its sub-millimolar binding affinity at site I [18], Mg^2+^ and Ca^2+^ are considerably more likely to be bound (see Fig. 7). Meanwhile, water frequently exchanged with K^+^ in site II, which is suggestive of a low affinity for the cation.

Based on our comparison with proteins that selectively bind K^+^, such as the KcsA K^+^ channel, selective binding of K^+^ over competing Ca^2+^ might best be afforded through placement of carbonyl oxygens at sufficiently large distances to accommodate the K^+^ ion’s larger volume. Namely, in K^+^ channels, oxygens from the backbone or side chains of amino acid forming the selectivity filter are exquisitely arranged to achieve precise pore size control and K^+^-oxygen interaction strength [52]. In contrast, it is likely that K^+^ exerts its agonistic effects on SERCA through binding the cytoplasmic domain, as was evidenced in a crystal structure determined by Sorenson et al, based on a bound K^+^ ion in the P-domain [5]. According to the K^+^-oxygen coordination pattern shown in Fig. S10, the MSA predicted potential for the P domain K^+^ is −2.38 kcal/mol, which is comparable to the values predicted for K^+^ bound to site I (−2.35 kcal/-mol). Consistent with this structural evidence, the E2P.2Ca dephosphorylation data indicate that K^+^ stimulates the Ca^2+^-release step in this state, whereby the lumenal Ca^2+^ affinity is reduced, rates of exchanging lumenal Ca^2+^ with lumenal solvated Ca^2+^ are accelerated [53], and Ca^2+^ release is enhanced [54]. Similarly, dephosphorylation of the E2P state is blunted in the absence of K^+^ [47].

Although the thermodynamics of K^+^ binding are unfavorable relative to Mg^2+^ and Ca^2+^, there remains the possible role of K^+^ in shaping the kinetics of SERCA function. It has been speculated, for instance, that K^+^ accelerates Ca^2+^ binding by first transiently occupying site I, after which it exchanges quickly with a Ca^2+^ at site II [54]. K^+^-facilitated exchange could therefore permit faster incorporation of Ca^2+^ into site I, as opposed to the direct migration of Ca^2+^ into a site that is potentially only partially-formed [54]. This interpretation is consistent with our observations from molecular dynamics simulations, and was initially reported in simulations of Ca^2+^-free SERCA by Musgaard et al [55].

### 3.2 Relating cation affinity to ATPase activity

In our opinion, the bridging of molecular-level simulation and MSA thermodynamic data with state-based models represents a significant contribution toward multi-scale modeling of steady-state SERCA activity. Numerical and computational modeling of SERCA activity has spanned phenomenological frameworks, such as Hill-type models [56–58] to those representing distinct stages of the catalytic process as states [41, 42, 59–61]. Our implementation is based on the stepwise binding of Ca^2+^ originally proposed by Inesi et al [41], but additionally considers Mg^2+^- and K^+^-bound states, followed by a reduction scheme to lump the E1 states separately from those comprising the E2 configurations. Significantly, the states defined in our model coincide with SERCA structures determined by x-ray crystallography and are ordered in a manner consistent with assays of SERCA function [62] In contrast, the state model proposed by Tran et al [42] assumed that ATP binding precedes the binding of two Ca^2+^ ions, which has not been experimentally verified.

Existing models of SERCA function have faithfully recapitulated the pump’s activity. However, an advantage of our state-based model is that its alignment with experimentally-determined structures permits us to directly investigate how structural modifications might impact activity. For example, we performed MSA/MD simulations of two mutated SERCA variants that yielded modified Ca^2+^ binding constants that we used to predict SERCA activity (see Fig. 8).While we acknowledge that there are likely myriad changes to the pump’s energetics following mutation that could be accounted for in the state-based model, our implementation here is a significant step toward structure-based modeling of SERCA activity. At a minimum, our fitted state model is consistent with steady-state pump rates data collected as a function of Ca^2+^ by Cantilina et al [41], Ca^2+^-saturated versus Mg^2+^ [17], as well as molecular simulations that predict Mg^2+^ inhibition of the pump [21]. Furthermore, by considering different linkages of K^+^- and Mg^2+^-bound states to the reaction scheme, we were able to determine that assumptions of Mg^2+^ inhibition and K^+^ agonist were most consistent with experimental data collected by Guillain et al [17]. In principle, multiscale models of SERCA activity that include structure-derived thermodynamic information could permit in silico investigations of how disease-associated SERCA mutations [63, 64], post-translational modifications [65], and binding of regulatory proteins such as phospholamban [59] affect pump function.

### 3.3 Limitations

There are several limitations of our approach that could be addressed in future studies. Our study largely focused on conformational changes and energetics of the cations’ immediate coordination sphere. Mg^2+^ and Ca^2+^-bound structures (PDB ID: 1su4 and 3w5b, respectively) deposited in the Protein Data Bank indicate substantial differences in the conformations of the transmembrane bundle helices and cytosolic domains that will necessarily contribute to the free energies of binding. For this reason, the free energy differences implied in experimental measurements of cation binding affinities reflect contributions from both ion coordination and the pump’s different conformations. It is also important to emphasize that the MSA assumptions of a spherical binding volume within which amino acid oxygens are immobilized crudely approximates the actual binding site. Thus, the predicted thermodynamic quantities are most appropriate for rank-ordering different ion/binding site configurations. Additionally, there is evidence [16, 62] that Ca^2+^ binding is cooperative, although here we treat the binding events independently. It may be possible to partially recover some of the cooperative behavior by modeling SERCA with only one bound Ca^2+^ and using end-point methods such as Molecular Mechanics/Poisson Boltzmann Surface Area (MM/PBSA) [66] to estimate energy changes upon binding a second Ca^2+^. To our knowledge, however, a half-saturated Ca^2+^-bound SERCA structure has not been determined, which would challenge efforts to validate predictions. Additionally, in the sequential binding model used by Inesi and others [6, 41] (*E*1 → *E*1.*Ca* → *E*1′*Ca* → *E*1′2*Ca*), the apparent kinetics governing the transition between the E1.Ca to E1’Ca states can vary depending on a variety of factors, including the presence the regulatory protein phospholamban [59, 67]. Thus, extending this model to broader conditions and regulatory proteins would require careful consideration of conformational changes that might accompany the E1.Ca to E1’Ca transition. Lastly, the Ca^2+^-binding steps in our model are assumed to be in steady state relative to the substantially slower transitions between the E1 and E2 states. In the event that the pump is subject to rapid Ca^2+^ oscillations, such as spontaneous Ca^2+^ spiking in glia [68], the steady-state approximation may be unreliable.

## 4 CONCLUSIONS

We utilized molecular dynamics simulations, mean sphere approximation theory and state-based modeling to probe effects of Ca^2+^, Mg^2+^ and K^+^ binding on the SERCA pump cycle. The MD and MSA approaches indicate that favorable binding of Ca^2+^ in the wild-type SERCA configuration is facilitated through a high degree of coordination by amino acids comprising the binding sites, as well as significant contributions from water coordination. This coordination pattern appears to be impaired in the E309Q and N796A variants; using MSA theory, we found that the chemical potential of Ca^2+^ binding is less favourable relative to wild-type as a result. Hence, mutations near the Ca^2+^ binding domains that alter the coordination number, hydration and binding site volume can be expected to modulate Ca^2+^ affinity in a manner qualitatively described by the MSA theory. Similarly, the coordination patterns exhibited in the Mg^2+^ and K^+^-bound structures led to less favorable binding estimates from MSA-theory. These findings were qualitatively consistent with measured affinity data reported in the literature [15–18, 45, 54]. Additionally, we developed a state-based model of SERCA activity that we used to: 1) relate Ca^2+^ binding affinities to the SERCA pump rate; and 2) determine whether Mg^2+^ and K^+^ are pump agonists or inhibitors. We found that the state model treating Mg^2+^ as an inhibitor and K^+^ as an agonist (Mg_−_K_+_) was most consistent with experimental data. Despite the limitations of the assumptions used for the various models, we believe that the approach provides an attractive framework for evaluating allosteric functional effects of ion binding on SERCA, which may be extendable to other Ca^2+^ transporters, such as P2X4 [69].

## 5 METHODS

### 5.1 Construction of the SERCA systems

Molecular dynamics (MD) simulations on wild-type (WT) SERCA and two variants of SERCA: E309Q and N796A were performed; the latter variants were chosen given dimished site II Ca^2+^-binding [44]. For the WT SERCA system, we considered apo (free of bound ions) Ca^2+^-bound, Mg^2+^-bound and K^+-^bound states. Ca^2+^-bound (PDB ID: 1su4 [3]) and Mg^2+^-bound (E1.Mg, PDB ID: 3w5b [37]) SERCA X-ray crystal structures were used as the starting structures. The apo SERCA system was constructed based on 1su4 with two Ca^2+^ ions removed as done in [70]. The structure of the site I or II bound K^+^ SERCA has not been determined, thus we created the structure based on the Mg^2+-^bound variant. For the N796A mutant, Ca^2+^-bound and Mg^2+^-bound cases were considered, based on mutating N796 to alanine. Similarly, for the E309Q mutant, we obtained two rotamers compatible with the binding site as evaluated through UCSF Chimera [71]. These E309Q rotamers were designated as “E309Q r1” and “E309Q r2”. All SERCA cases considered in present study are summarized in Table S1.

The cation coordinating acidic residues E309, E771 and D800 were assumed to be deprotonated, while E908 was protonated, consistent with [21]; further a disulfide bond was introduced between residues C876 and C888. The system was inserted into a homogeneous 1-palmitoyl-2-oleoyl-*sn*-glycero-3-phosphocholine (POPC) lipid bilayer of POPC via the Membrane Builder [72] within the CHARMMGUI software [73]. This system was solvated via TIP3P waters [74] using a 20 Å margin perpendicular to the membrane. Counterions K^+^ and chloride (Cl^−^) were added into the system via Monte-Carlo method to neutralize the system and maintain an ionic strength of 0.15 M. Both the solvation and ion-adding were performed via the Solvator module within CHARMM-GUI online-server. The final system contained ~255,000 atoms including lipid bilayer with 461 lipids, and ~59,000 TIP3P water molecules and was parameterized by the CHARMM36 force field [75, 76].

### 5.2 Molecular dynamics simulations

MD simulations were performed using NAMD [77]. The system was first subjected to an energy minimization process consisting of 2000 steps’ steepest descent (SD) and 2000 steps’ adopted basis Newton rRphson (ABNR) algorithm. For each case, the minimized system was heated from 0 K to 303.14 K over 25 ps with 1 fs timestep via the Langevin thermostat, using randomized velocities for three triplicate preparations. For each replica, harmonic constraints were introduced during minimization and heating on protein side chains, protein backbone atoms, lipid heavy atoms and ions. The force constants of constraints were set to 5, 10, 10 and 1.0 × 10^1^ 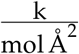, respectively, and were gradually reduced to zero over five equilibration steps of 20 ps in duration. 100 ns production simulations was then performed on the equilibrated system with the Shake algorithm [78], using 2 fs timesteps.

### 5.3 Simulation trajectory analysis

Standard trajectory analyses including RMSF and coordination oxygen/Ca^2+^ distances were computed using Lightweight Object-Oriented Structure Library (LOOS) [79]. Coordination pattern analysis on each cation with oxygen atoms from nearby amino acids were performed in each binding site. This consisted of extracting residues within 20 Å from D800 (the shared residue of the conventional two Ca^2+^-binding sites in SERCA) based on the Ca^2+^-bound SERCA crystal structure (PDB ID: 1su4). The water density around cation was computed via the radial command implemented in CPPTRAJ [80] and analyzed via GIST (as described below).

The MSA is a mean-field model which estimates cation chemical potentials in electrolyte solution with finite-sized ions. In this study, the SERCA cation binding sites were treated as confined filters filled with oxygens from coordinating residues and water molecules. The MSA model calculates cation distributions between bulk electrolyte solution that minimize the chemical potential for partitioning ions into a finite size volume occupied by coordinating oxygens; these volume and number of oxygens were determined by MD as in Kucharski et al[34]. The free energy expression in this model was assuming negative-charged spherical domains was based on a chemical potential accounting for electrostatic and hard-sphere interactions between ions inclusive of oxygens, as described in [35, 81, 82]. In this representation, which is analogous to the Ca^2+^ binding domain in *β*-PV and calcium channels [33–35], ionic species that have a negative chemical potential in the binding filter are thermodynamically favored to bind. We include in this approach a solvation contribution as estimated via generalized Born theory of ion hydration energies:

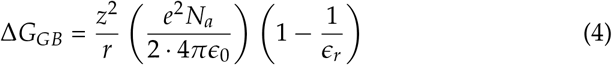

where *z* is charge number, *r* is radii, *e* is electron charge, *N*_*a*_ is the Avogadro constant, *ϵ*_0_ and *ϵ*_*r*_ are vacuum dielectric and the relative dielectric constant of the solvent. Additional details are elaborated in the supplementary material of [34].

In the SERCA system, the cation/SERCA configurations can present differing numbers of coordination oxygen atoms (see Fig. 4 and Table S2) and volumes, as shown previously for *β*-PV in [34]:

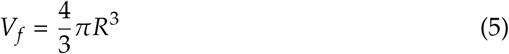

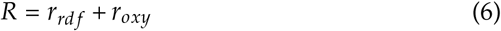

where *r*_*rdf*_ is the radii of optimal coordination sphere which can be obtained from the cation-oxygen coordination pattern analysis based on MD simulations (see Fig. 4) while *r*_*oxy*_ is the radii of oxygen atom. Charges and radii values of all ions are taken from the Li-Merz work [83]. The specific parameters used in the MSA calculations are listed in Table S2.

### 5.4 State-based model of the SERCA pump cycle

SERCA pumping is characterized by two prominent states, E1 and E2, comprised of microstates corresponding to various conformations or bound states of the pump (Fig. S4). In the E1 state, two Ca^2+^ ions bind to SERCA through cooperative mechanism followed by the binding of MgATP [41]. We represent each state as by an ordinary differential equation of the form

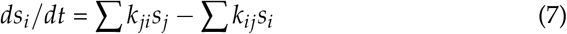

where *s*_*i*_ is state i, and *k*_*ij*_ describe rates for transitioning between states i and j. The models tested in this paper are summarized in Fig. S4. While the cooperative Ca^2+^ binding mechanism in the E1 state is well-accepted, the Ca^2+^ release mechanism of the E2 state is not clearly resolved, therefore we use the state representations proposed in Tran et al [42].

Based on the technique proposed by Smith et al [84], we can applied rapid equilibrium assumption to fast processes (those within E1 or E2) relative to the slow rates of transitions between the E1 and E2 states to simplify the model. As shown in Fig. S4, states in dashed boxes were assumed to be in quasi-steady-state and thus were lumped together. Hence, the original multi-state models were simplified into a two-state model (see bottom Fig. S4) in which new apparent rate constants were derived based on original rate constants. The resulting equations are summarized in the supplement Sect. S1.3.

For the lumped 2-state model, the turn over rate is derived as in Tran’s paper [42]:

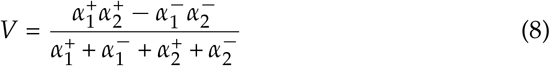

where 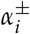 are apparent rate constants between lumped two states (the expressions are given in Sect. S1.3). The final parameters used are listed in Table S3.

## 6 ACKNOWLEDGEMENT

Research reported in this publication was supported by the Maximizing Investigators’ Research Award (MIRA) (R35) from the National Institute of General Medical Sciences (NIGMS) of the National Institutes of Health (NIH) under grant number R35GM124977, as well as the award 1F32HL114365-01A1. This work used the Extreme Science and Engineering Discovery Environment (XSEDE), which is supported by National Science Foundation grant number ACI-1548562. We thank Michael Autry for discussion of this work.

## S1 Supplementary Information (SI)

### S1.1 Supplementary Tables

**Table S1:** MD simulation cases

| Table S1: MD simulation cases |  |  |  |
| --- | --- | --- | --- |
| Cases | Cation occupancy | PDBID of ini. struc | Simulation length (ns) |
| WT | 2 Ca<sup>2+</sup>s | 1su4 | 300 |
| WT | 1 Mg<sup>2+</sup> | 3w5b | 300 |
| WT | 2 K<sup>+</sup>s | 3w5b | 300 |
| N796A | 2 Ca<sup>2+</sup>s | 1su4 | 400 |
| N796A | 1 Mg<sup>2+</sup>s | 3w5b | 355 |
| E309Q_r1<sup>a</sup> | 2 Ca<sup>2+</sup>s | 1su4 | 240 |
| E309Q_r2<sup>b</sup> | 2 Ca<sup>2+</sup>s | 1su4 | 170 |

<sup>a,b</sup>Different rotamers of Q309 side chain

### S1.2 Supplementary Figures

**Table S2:** Parameters used in SERCA MSA model. The diameter and charge of ions were adapt from Li-Merz work [83] and the solvation energy for ions were taken from table 1 in Nonner et al [33].

| Parameter | Definition | Values |
| --- | --- | --- |
| $N_{oxy}$ at $\text{Ca}^{2+}$ site I | no. of amino acid oxygen | 5.5 |
| $N_{oxy}$ at $\text{Ca}^{2+}$ site II | no. of amino acid oxygen | 6.7 |
| $N_{oxy}$ at $\text{K}^+$ site I | no. of amino acid oxygen | 3.9 |
| $N_{oxy}$ at $\text{Mg}^{2+}$ site | no. of amino acid oxygen | 4.2 |
| $r_{rdf}$ at $\text{Ca}^{2+}$ site I, II | optimal coordination radii | 2.5 Å |
| $r_{rdf}$ at $\text{Mg}^{2+}$ site | optimal coordination radii | 2.0 Å |
| $r_{rdf}$ at $\text{K}^+$ site I | optimal coordination radii | 2.9 Å |
| $\rho_i$ | Bath [KCl] | 150 mM |
| | bath [$\text{MgCl}_2$] | 1 mM |
| | bath [$\text{CaCl}_2$] | 0.1 nM to 0.1 M |
| $\epsilon_f$ | filter dielectric | 25.0 |
| $\epsilon_b$ | bath dielectric | 78.4 |
| $\sigma_O$ | O diameter | 0.354 nm |
| $\sigma_{Ca}$ | $\text{Ca}^{2+}$ diameter | 0.272 nm |
| $\sigma_{Mg}$ | $\text{Mg}^{2+}$ diameter | 0.236 nm |
| $\sigma_K$ | $\text{K}^+$ diameter | 0.352 nm |
| $\sigma_{Cl}$ | $\text{Cl}^-$ diameter | 0.454 nm |
| $z_O$ | O charge | -0.7469 e |
| $z_{Ca}$ | $\text{Ca}^{2+}$ charge | 1.77 e |
| $z_{Mg}$ | $\text{Mg}^{2+}$ charge | 1.69 e |
| $z_K$ | $\text{K}^+$ charge | 1 e |
| $z_{Cl}$ | $\text{Cl}^-$ charge | -1 e |
| $w_{Ca}$ | solvation energy | -648.65 kT [50] |
| $w_{Mg}$ | solvation energy | -779.94 kT [50] |
| $w_K$ | solvation energy | -141.59 kT [50] |
| $w_{Cl}$ | solvation energy | -122.63 kT [50] |
| $kT/zF$ | | 25.6 mV |
| $V_f$ of $\text{Ca}^{2+}$ sites | filter volume | 0.326 nm<sup>3</sup> |
| $V_f$ of $\text{Mg}^{2+}$ site | filter volume | 0.224 nm<sup>3</sup> |
| $V_f$ of $\text{K}^+$ site I | filter volume | 0.426 nm<sup>3</sup> |

**Table S3:** Parameters used SERCA state-based model models in present work.

| Parameter | Value | reference |
| --- | --- | --- |
| $K_{d,Mg}$ | $4.93 \times 10^{-5} \text{ M} / 5 \times 10^{-4} \text{ M}$ | MSA/Expt. |
| $K_{d,K}$ | $3.3 \times 10^{-3} \text{ M} / 9 \times 10^{-3} \text{ M}$ | MSA/Expt. |
| $K_{d,2}$ | $2.24 \times 10^{-3} \text{ M}$ | [42] |
| Ca(SR) | $1 \times 10^{-6} \text{ M}$ | [41] |
| [MgATP] | $3 \times 10^{-4} \text{ M}$ | [41] |
| [MgADP] | $1 \times 10^{-6} \text{ M}$ | [41] |
| [Pi] | $1 \times 10^{-6} \text{ M}$ | [41] |
| $k_1^+$ | $1 \times 10^8 \frac{1}{\text{s}}$ | [41] |
| $k_1^-$ | $4 \times 10^2 \frac{1}{\text{s}}$ | [41] |
| $k_2^+$ | $3.0 \times 10^1 \frac{1}{\text{s}}$ | [41] |
| $k_2^-$ | $4.0 \times 10^1 \frac{1}{\text{s}}$ | [41] |
| $k_3^+$ | $4 \times 10^8 \frac{1}{\text{s}}$ | [41] |
| $k_3^-$ | $1.6 \times 10^1 \frac{1}{\text{s}}$ | [41] |
| $k_4^+$ | $2 \times 10^7 \frac{1}{\text{s}}$ | [41] |
| $k_4^-$ | $4.0 \times 10^1 \frac{1}{\text{s}}$ | [41] |
| $k_5^+$ | $1.5 \times 10^1 \frac{1}{\text{s}}$ | fitted |
| $k_5^-$ | $7.5 \frac{1}{\text{s}}$ | fitted |
| $k_6^+$ | $7.5 \frac{1}{\text{s}}$ | fitted |
| $k_6^-$ | $7 \times 10^9 \frac{1}{\text{s}}$ | fitted |

**Table S4:** Experimental dissociation constant of cations with SERCA

| Cation | Dissociation constant ($K_d$) | reference |
| --- | --- | --- |
| Ca<sup>2+</sup> site I | 0.018 - 0.023 $\mu\text{M}$ | [16] |
| Ca<sup>2+</sup> site II | 6.67 - 11.1 $\mu\text{M}$ | [16] |
| Mg<sup>2+</sup> | 5 mM / 0.5 mM<sup>a</sup> | [17] and [54] |
| K<sup>+</sup> site I | 0.625 mM | [18] |
| K<sup>+</sup> site II | 16.67 mM | [18] |
| K<sup>+</sup> binds to P-domain<sup>b</sup> | na | [5] |
| N796A and E309Q Ca<sup>2+</sup> site I | 0.5 $\mu\text{M}$ | [45] |
| N796A and E309Q Ca<sup>2+</sup> site II | no binding | [45] |
| E309Q with 2 Ca<sup>2+</sup> binding | 0.46 - 4.38 $\mu\text{M}$<sup>c</sup> | [15] |

<sup>a</sup> in [54] it was postulated that Mg<sup>2+</sup> binds to site II with $K_d = 0.5 \text{ mM}$ and this bound-Mg<sup>2+</sup> would exchange quickly with Ca<sup>2+</sup> and the consequence is Mg<sup>2+</sup> accelerates Ca<sup>2+</sup> bindings to SERCA; <sup>b</sup> a K<sup>+</sup> binding site located in P-domain was identified via X-ray crystallography; <sup>c</sup> This data is apparent affinity for both site I and II obtained by fitting to experimental data using Hill equation.

**Figure S1:**
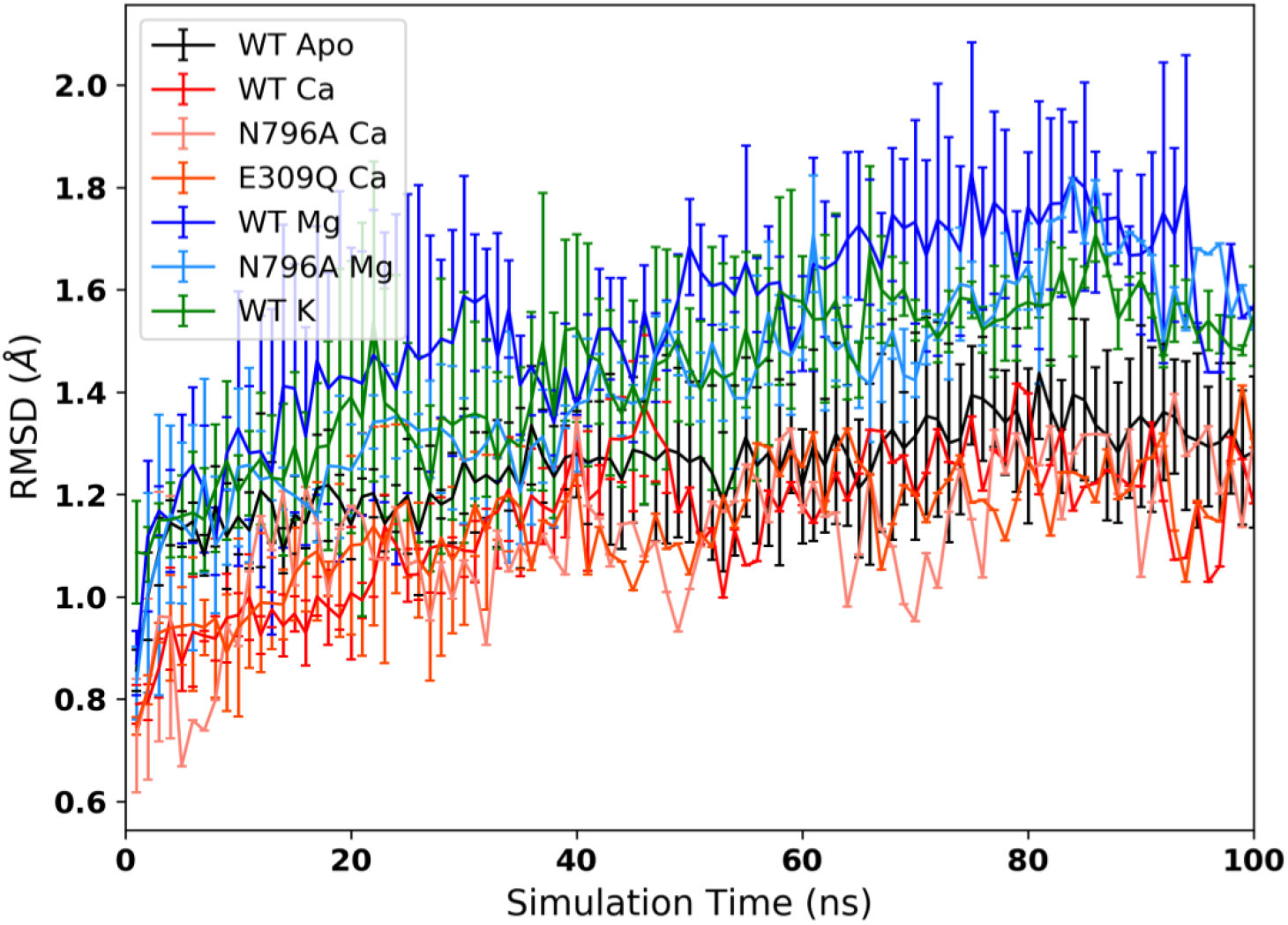
Back-bone atom RMSD of TM helices vs. simulations time. The error bars are calculated from the triplicate trajectories for each case.

**Figure S2:**
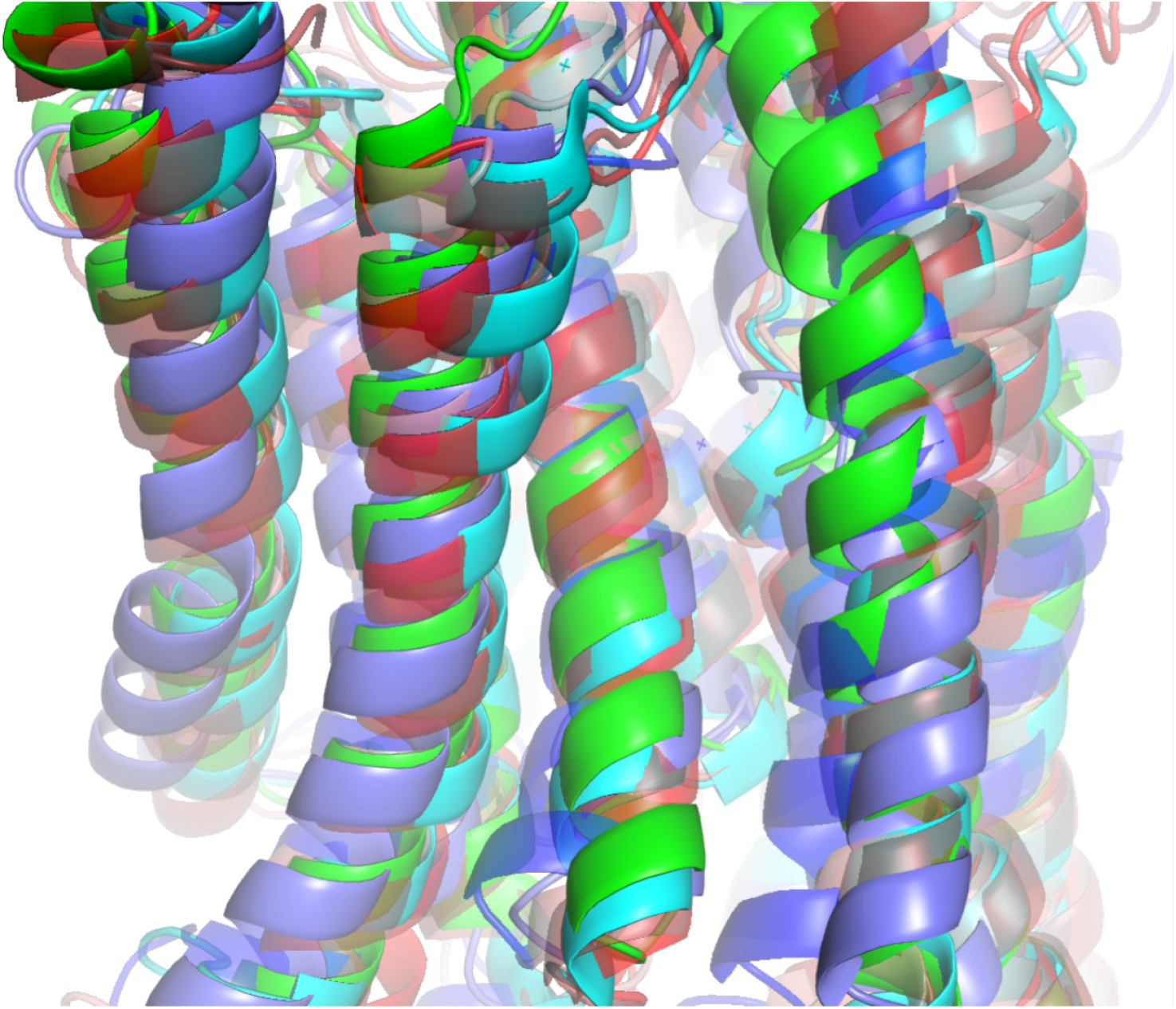
TM alignments of MD-simulated structures to x-ray crystal structures of PDB 1su4. The MD-simulated structures are extracted from MD trajectories at 80 ns and colored as red for Ca^2+^-bound cases, blue for Mg^2+^-bound cases and green for K^+^-bound cases. The crystal structure is colored in cyan.

**Figure S3:**
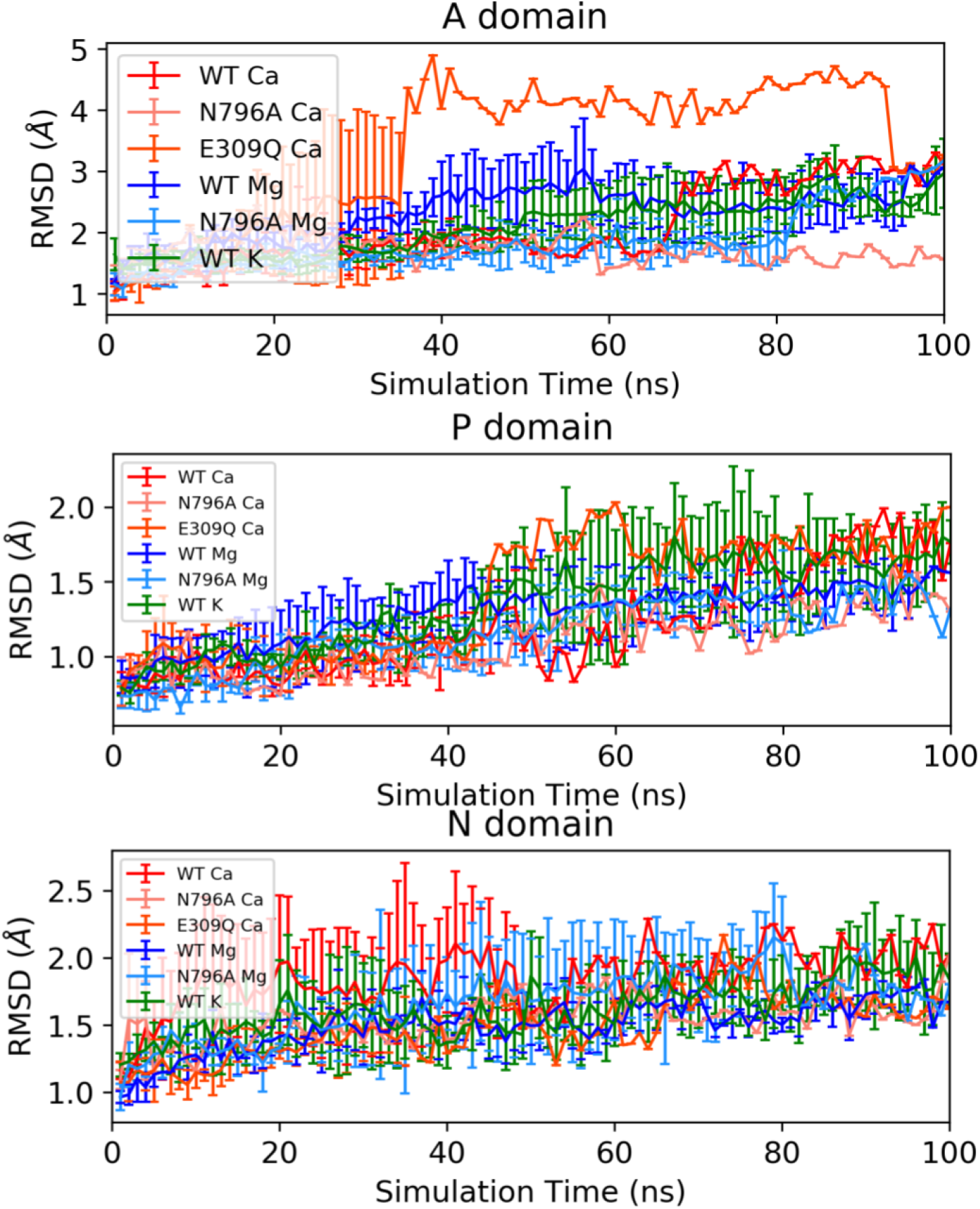
Back-bone atom RMSD of cytosolic domains vs. simulations time. Alignments were conducted using backbone atoms of cytosolic domains. The error bars are calculated from the triplicate trajectories for each case.

**Figure S4:**
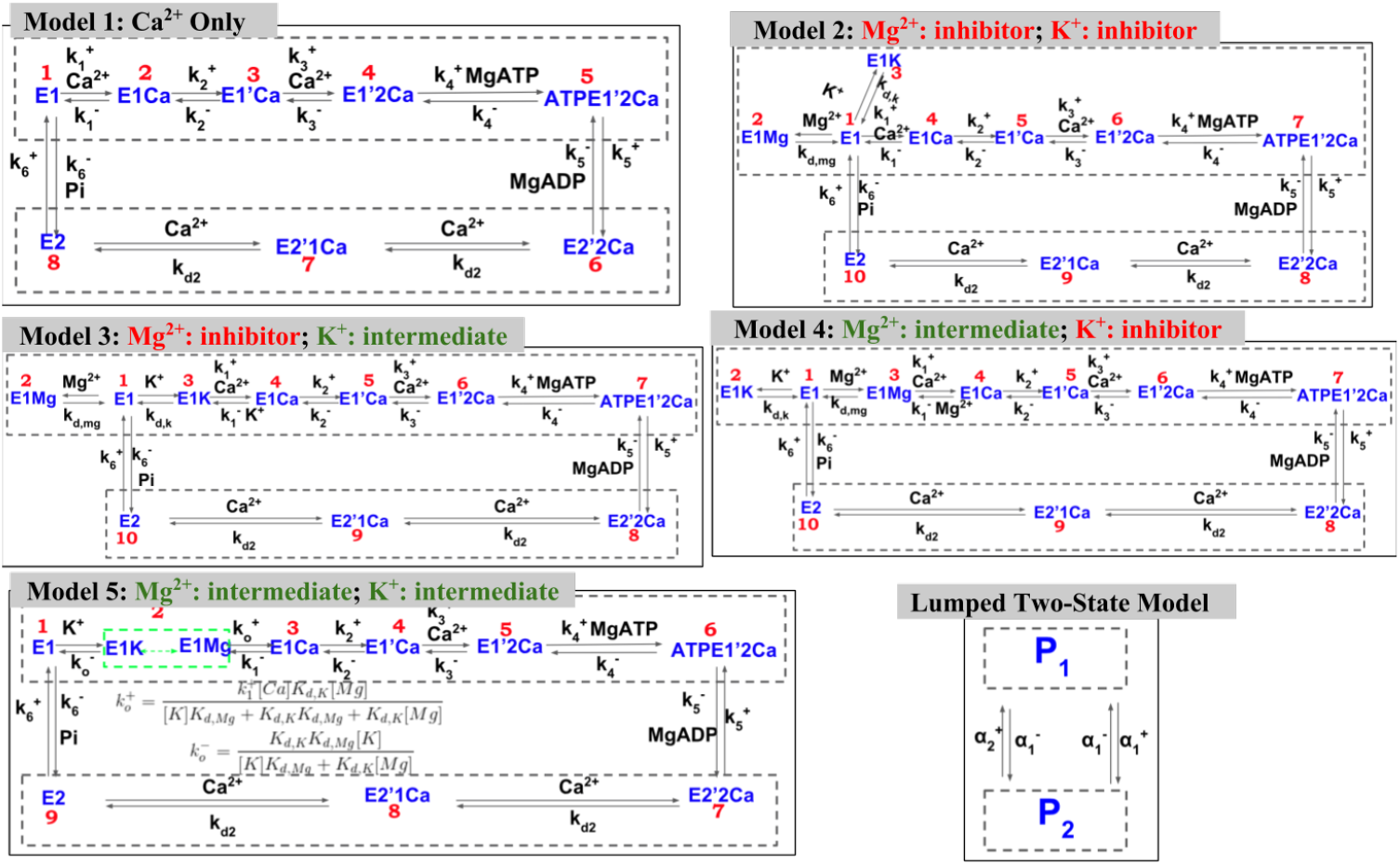
Schema of the five state-based model models of SERCA pump considered in present study. In Model 1 (Ca_*only*_), only the native Ca^2+^ cation is involved in the pump cycle. In Model 2 (Mg_−_K_−_), both Mg^2+^ and K^+^ are assumed to act as inhibitors of SERCA. In Model 3 (Mg_−_K_+_), Mg^2+^ is assumed to act as inhibitors of SERCA while K^+^ acts as intermediate. In Model 4 (Mg_+_K_−_), the roles of Mg^2+^ and K^+^ are opposite as Model 3, namely, Mg^2+^ is assumed to act as intermediate of SERCA while K^+^ acts as inhibitor. In Model 5 (Mg_+_K_+_), both Mg^2+^ and K^+^ are assumed to act as intermediates of SERCA. For all models, the states in dashed-box are lumped together based on the lumping strategy proposed by Smith et al [84] to form a two-state model. The apparent rates between the two-state model are given in supplementary material.

**Figure S5:**
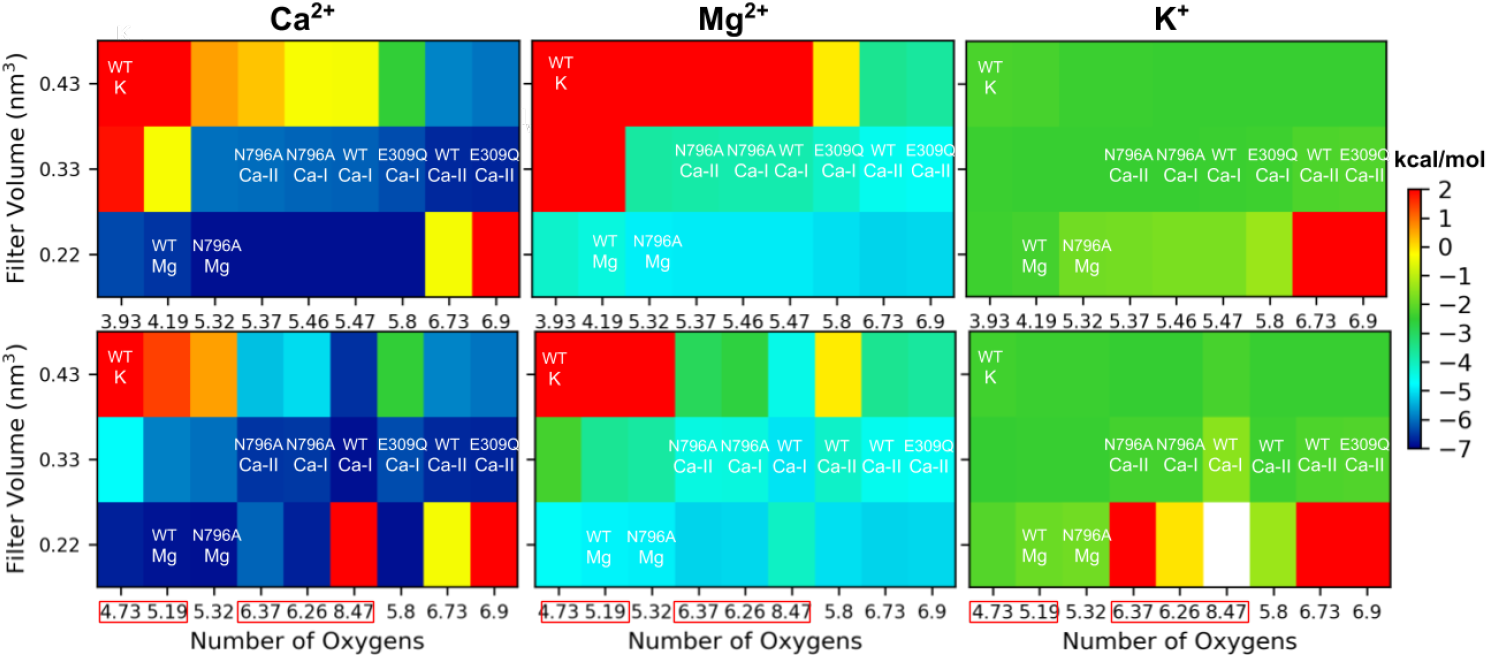
Chemical potential of cation in the filter calculated via MSA at various values of filter volume (*V*_*f*_) and coordinated oxygens. The upper and lower row represent results where coordinating water molecules were excluded and included, respectively. The red boxed xticks in lower row highlighted the total number of coordinating oxygens when water were take into account. The bath ionic strength are [KCl] = 150 mM, [MgCl_2_] = 2 mM and [Ca^2+^]= 1 *μ*M. The three filter volumes of the *y*-axis are optimal volumes for Ca^2+^ (0.33 nm^3^), Mg^2+^ (0.22 nm^3^) and K^+^ (0.43 nm^3^). The values at *x*-axis correspond to the optimal coordinating oxygens determined from MD simulations.

**Figure S6:**
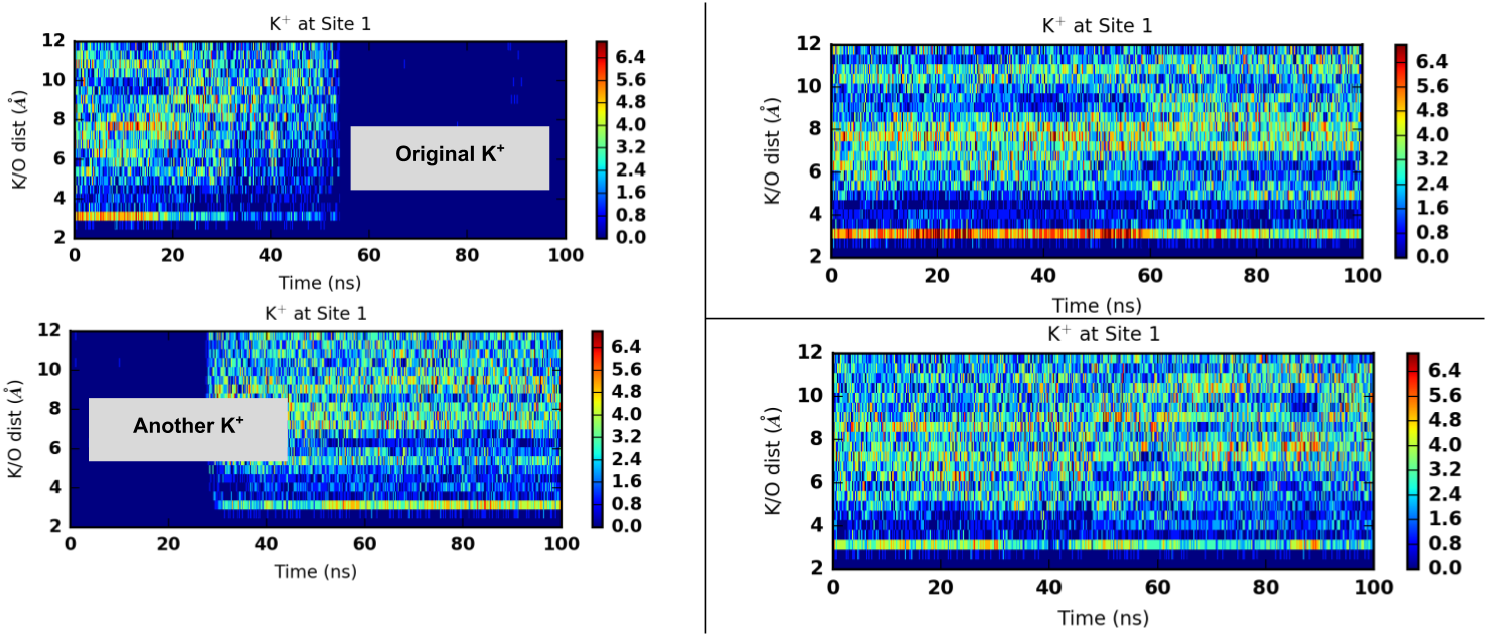
Coordination pattern of K^+^ at SERCA site I with residue oxygen atoms in the 1st 100 ns WT simulations. In the 1st 100 ns simulation, the original K^+^ at site I flee away after about 40 ns, however, another K^+^ will bind at site I afterwards. The radial distribution is calculated based on the second K^+^ after 40 ns.

**Figure S7:**
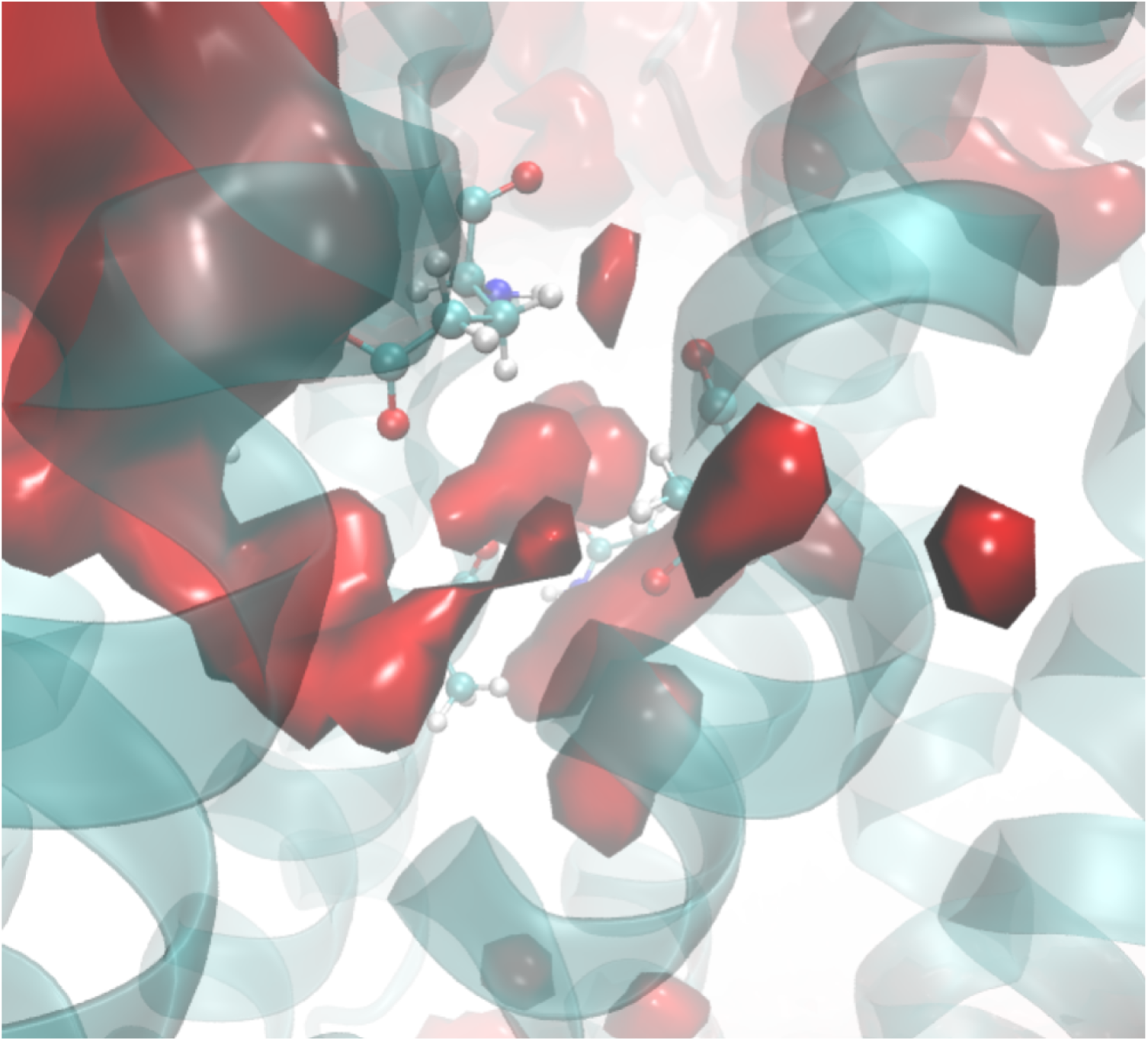
Water density in Ca^2+^ binding sites calculated from apo SERCA MD simulations via the VolMap module of VMD. The key coordinating residues are shown as sticks. The shown red isosurface depicts water density as 0.35.

**Figure S8:**
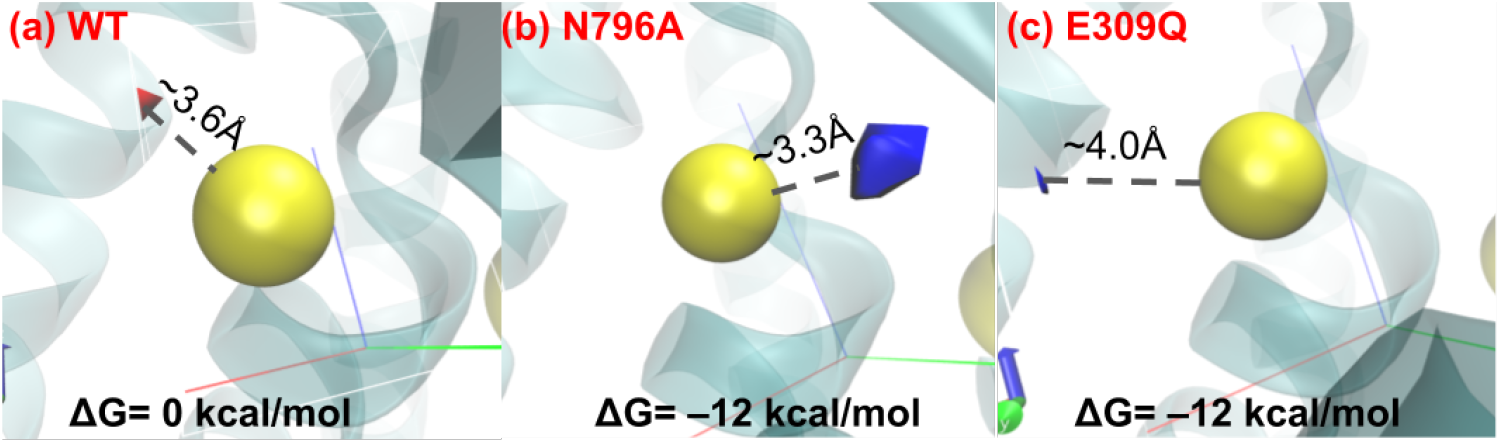
Free energy of water sites around site II Ca^2+^ calculated by GIST. For N796A and E309Q, the blue isosurface corresponds to water sites with Δ*G* = −12 kcal/mol. For WT the shown red isosurface depicts a water oxygen density as 0.075 while the Δ*G* is 0 everywhere over the grids. The distance between Ca^2+^ and isosurface is also shown.

**Figure S9:**
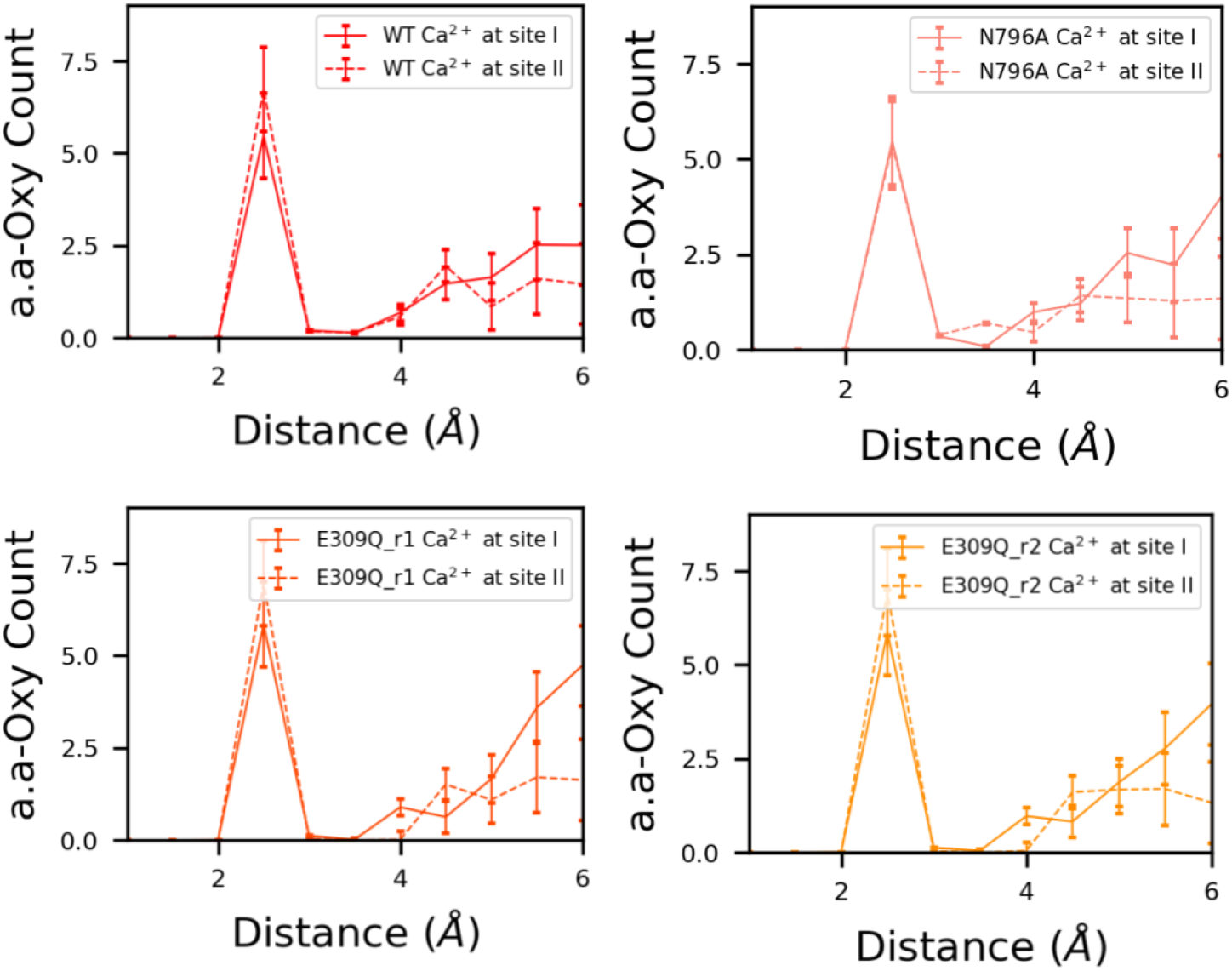
Radial distribution of oxygen around Ca^2+^ for each case.

**Figure S10:**
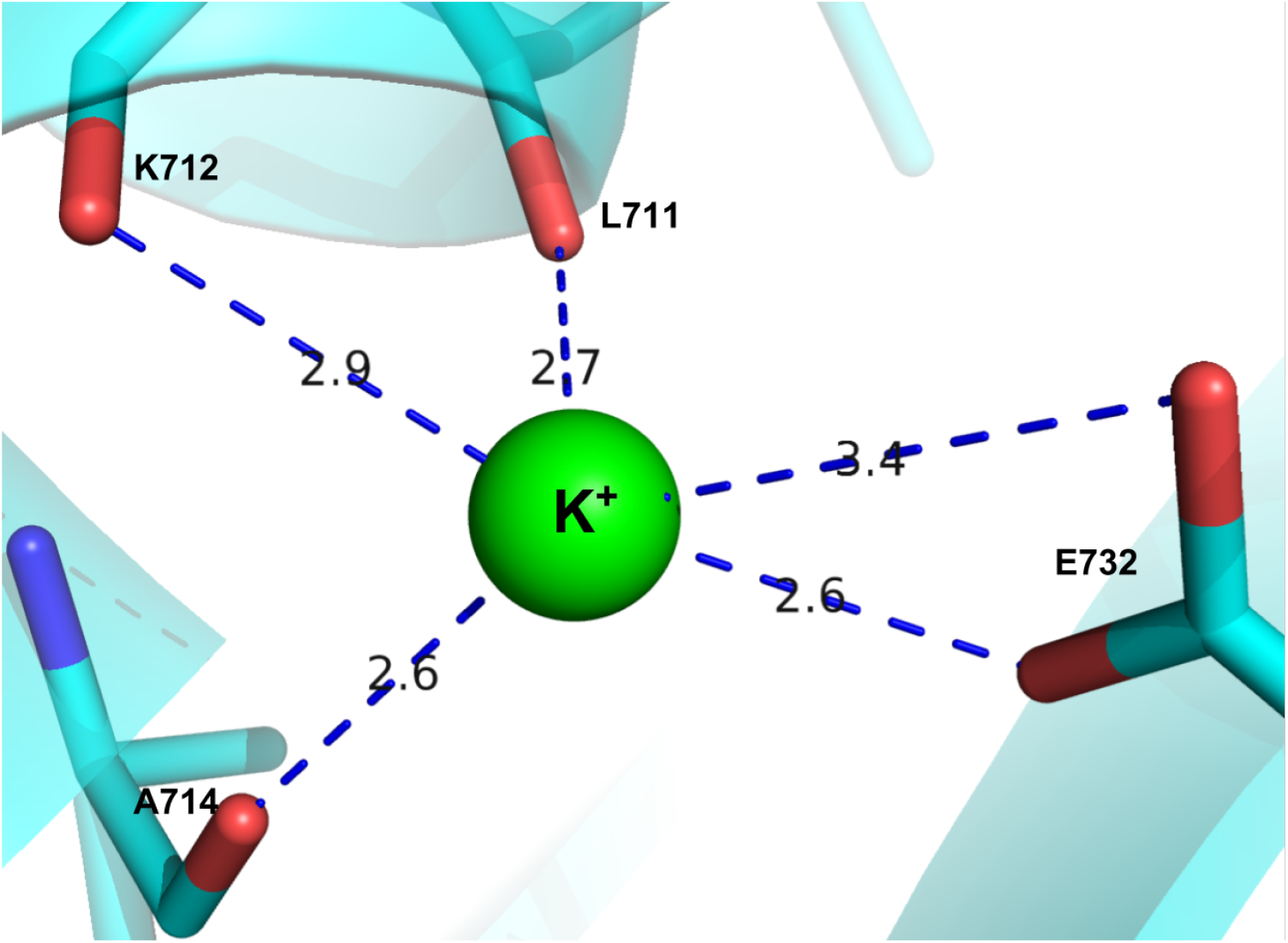
K^+^ binding site at P domain from crystal structure PDB 1t5s. Number of coordinating oxygens is five and filter volume is assumed to be 0.43 nm^3^.

**Figure S11:**
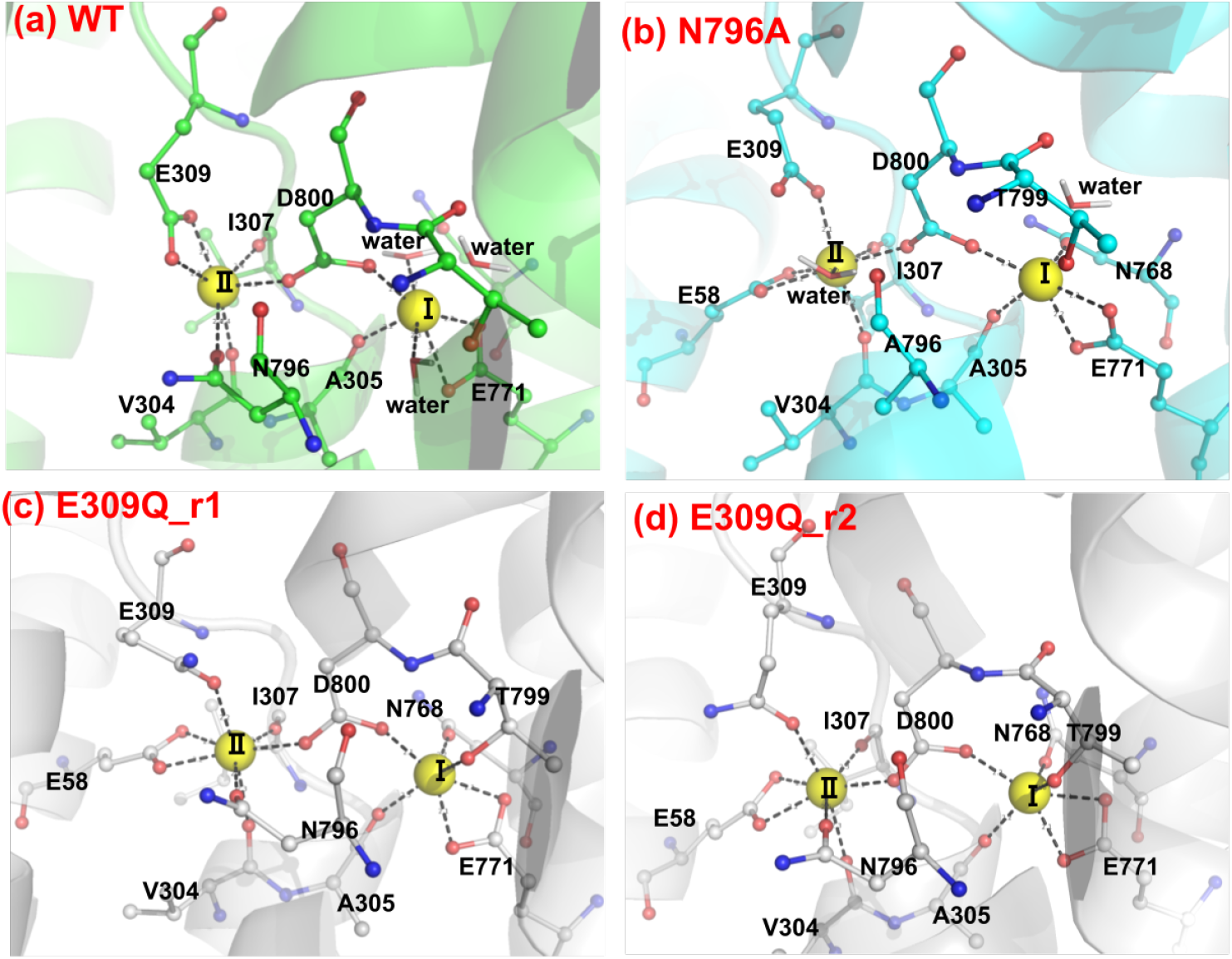
Representative coordination patterns of Ca^2+^ (yellow spheres) in the two binding sites from MD simulation in WT SERCA and mutants. Key residues are shown in sticks and balls. In WT site I, there are 3 waters coordinating with Ca^2+^, while in N796A/E309Q mutants, site I Ca^2+^ has one/zero coordinating water molecules.

**Figure S12:**
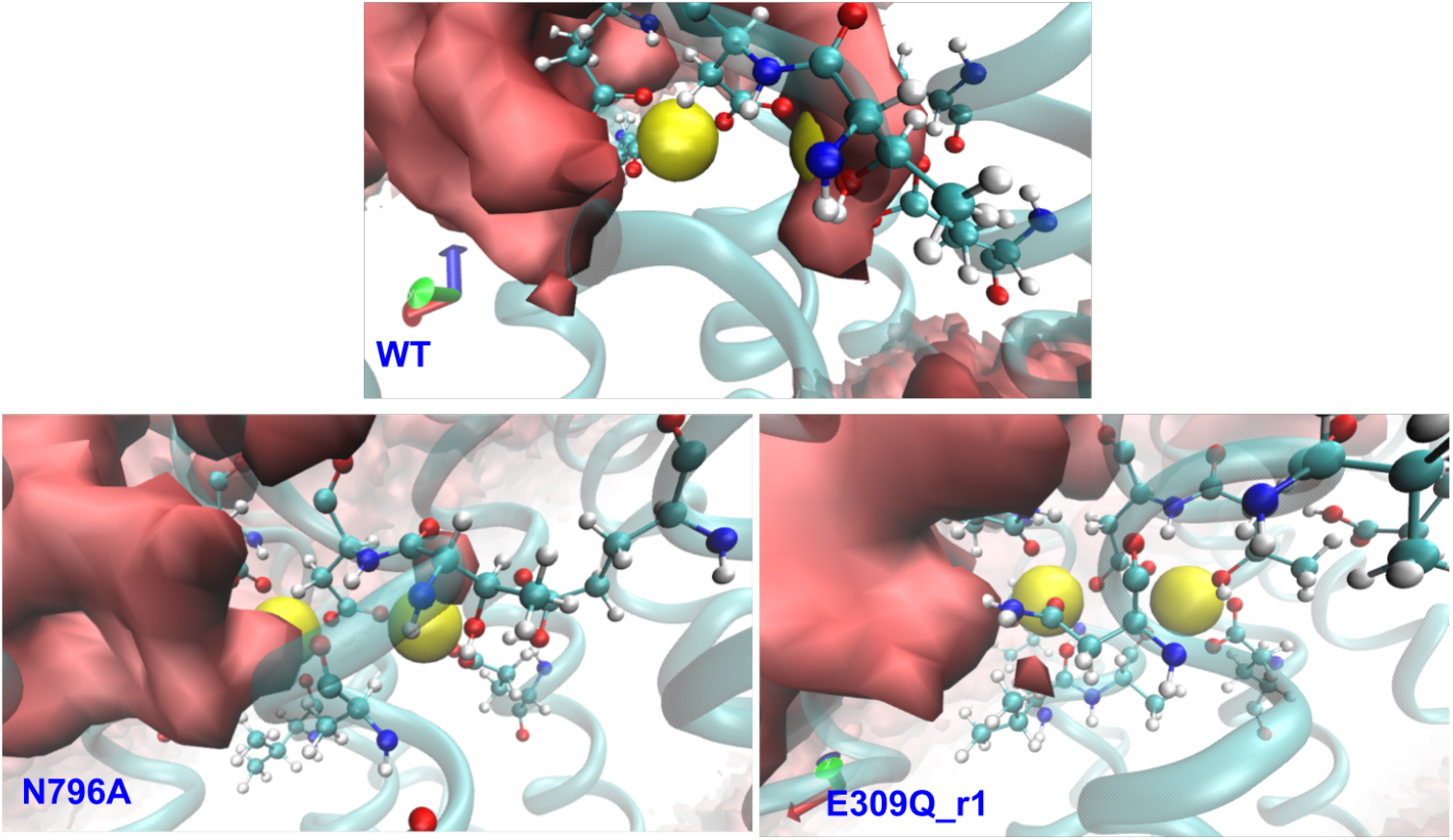
Water density around Ca^2+^ (yellow spheres) calculated from MD simulations via the VolMap module of VMD. The shown red isosurface depicts water density as 0.2.

**Figure S13:**
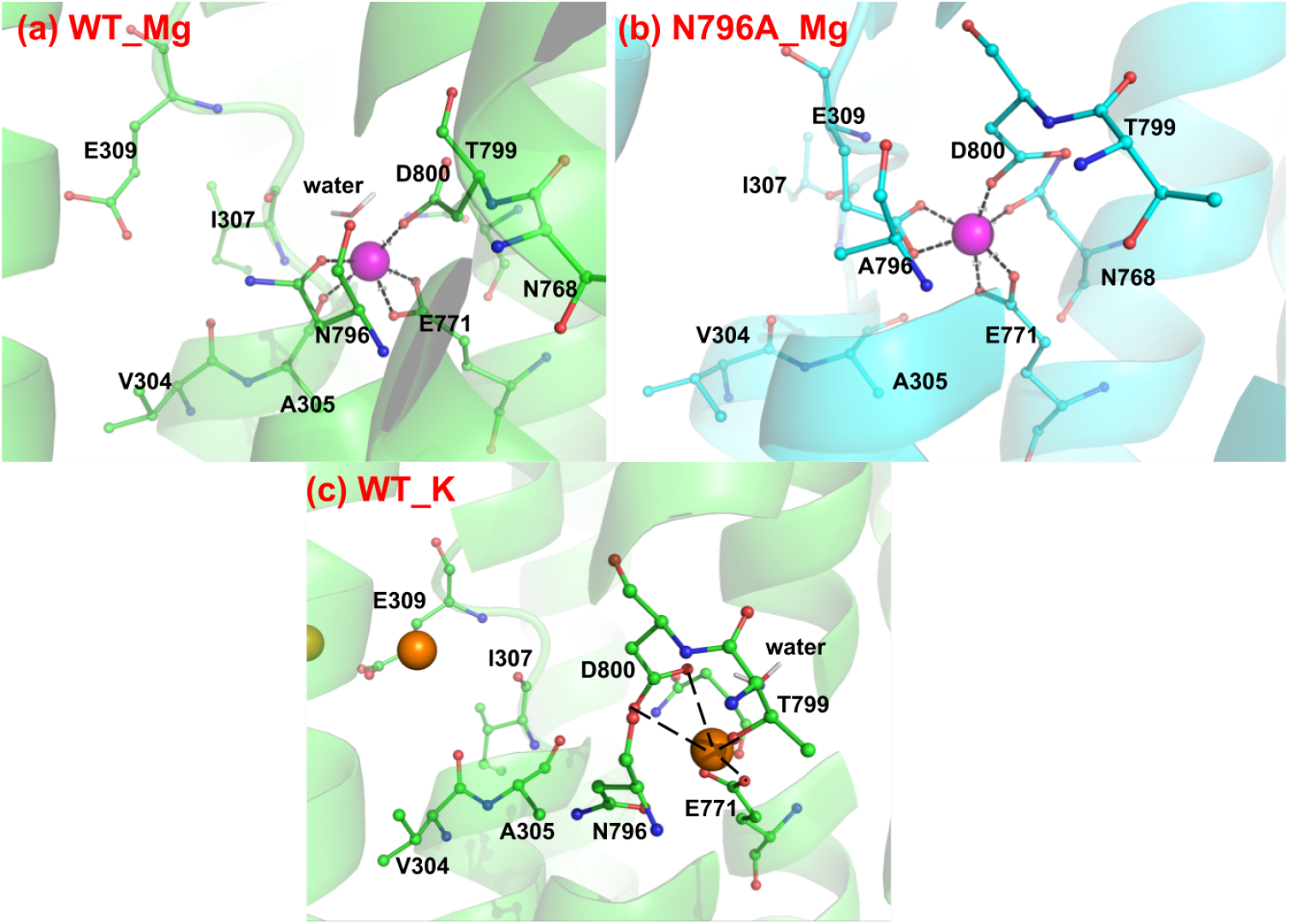
Representative coordination patterns of Mg^2+^ (magenta spheres) and K^+^ (orange spheres) in the binding sites from MD simulation in WT SERCA and mutants. In both WT and N796A, Mg^2+^ resides in a ‘hybrid’ binding site rather than the conventional two Ca^2+^ binding site. K^+^ at site I is stable and not stable K^+^ binds at site II.

### S1.3 Supplementary Methods

The Grid Inhomogeneous Solvation Method (GIST) model [85, 86] was used to characterize the theromodynamic properties of coordinating water. GIST calculations were performed via the SSTmap program [87]. Before GIST calculations, the MD trajectory was first subjected to rms fitting on key cation coordinating residues to align cation binding sites. The C*α* atoms of residues V304, I307, N768, E771, N796 and D800 were used for this rms fitting. The grid center required in GIST was defined by the cation coordinates obtained from the last frame in the rms-fitted trajectory. Grid dimensions were set to 16, 20 and 20 with grid spacing being 0.5 Å. The free energy of water at each grid point was calculated as Δ*G* = *E*_*sw*_ + *E*_*ww*_ − *dTS*_*orient*_ − *dTS*_*trans*_ where *E*_*sw*_ and *E*_*ww*_ are solute-water and water-water interaction energies while *dTS*_*orient*_ and *dTS*_*trans*_ are orientational and translational entropic energies.

Here we summarize our lumping strategy used to consolidate the SERCA model into fewer (macro) states, by assuming rapidly-exchanging ‘micro’ states are in pseudo-equilibrium. The basic approach follows that of Smith et al [84] as illustrated by the generalized example:

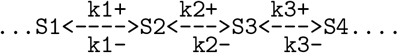

S1 to S4 are four states with forward and reverse rate constants connecting neighboring states. Suppose S2 and S3 are in rapid equilibrium with each other, namely 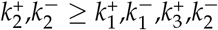. At any time:

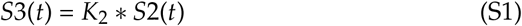

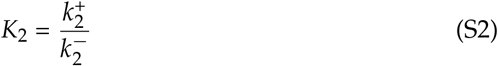

Now we lump S2 and S3 together to form a new pseudo state S23. For the lumped model, we will use new apparent rate constants 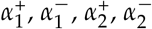 to represent the transition between states.

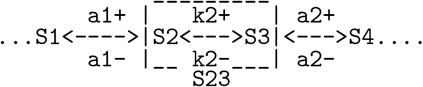

For the lumped model, we have:

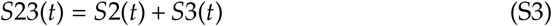

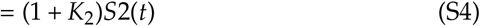

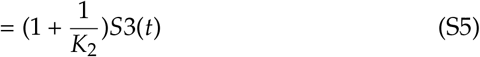

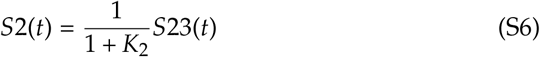

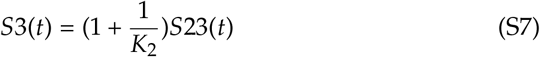

Now we consider relationship between original rate constants and the new apparent rate constants. In both models, for S4, we have:

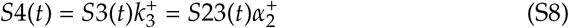

Now we put Eq(7) into Eq(8), we see clearly that:

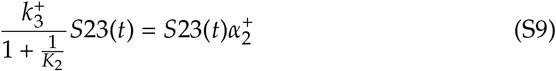

so we have the expression of 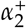 as:

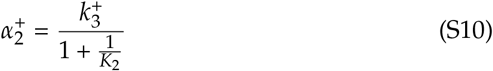

similarly, we can derive the expression of 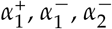:

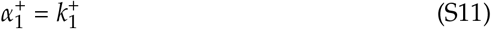

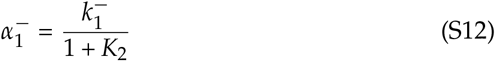

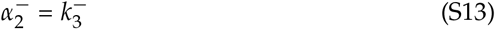

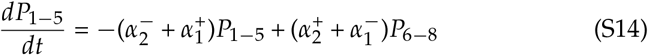

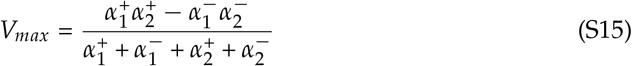

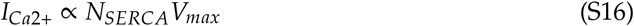

For Ca_*only*_ model, the apparent rate constants are:

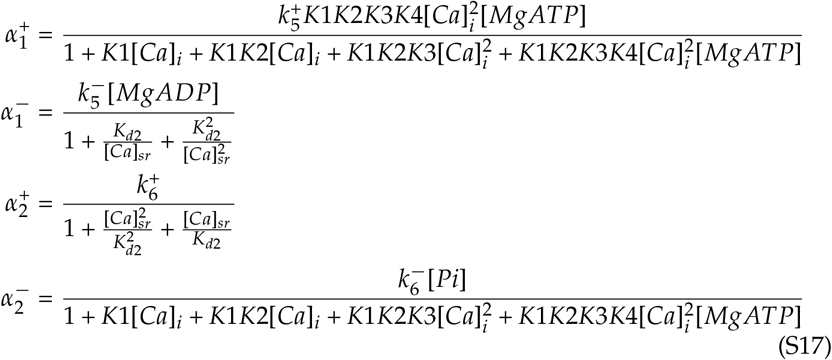

where 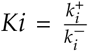 (*i* = 1, 2, 3, 4) and the value for each parameter is listed in Table S3. For models with Mg^2+^/K^+^ involved, the apparent rate constants can be expressed via the general expression Eq. S18:

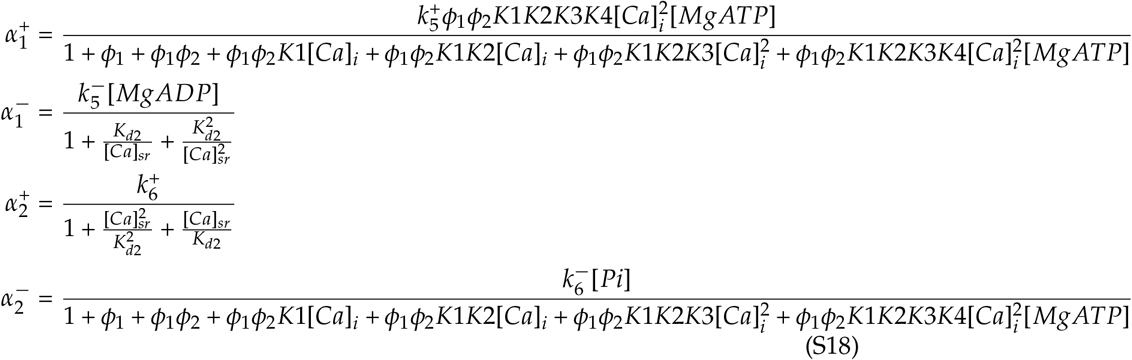

The expression of *ϕ*_1_ and *ϕ*_2_ differs in different models. For Mg_−_K_−_, we have:

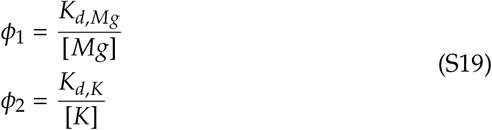

For Mg_−_K_+_, we have:

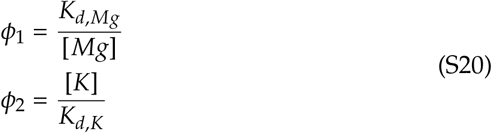

For Mg_+_K_−_, we have:

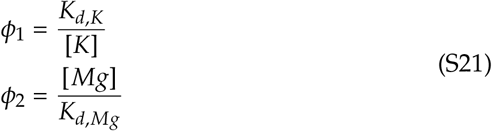

For Mg_+_K_+_, we have:

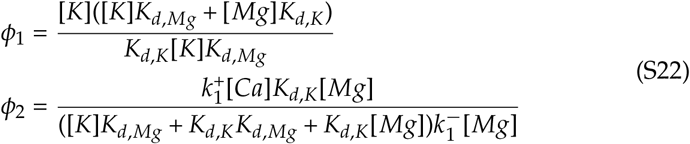

For state models with Mg^2+^ and K^+^ involved, to be consistent with experimental paper [17], we use the sum of E1.Mg and E1.2Ca state probaliby as a metric of Ca-binding, namely:

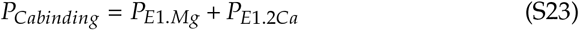

For Mg_−_K_+_, we have:

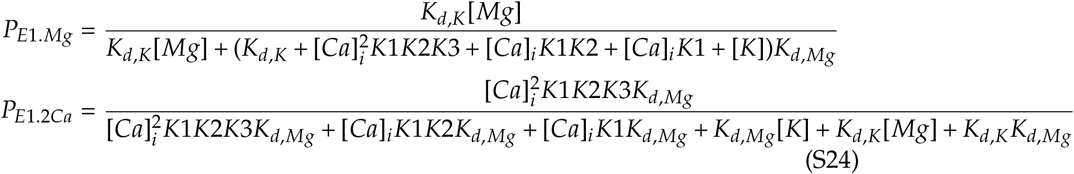

For Mg_+_K_−_, we have:

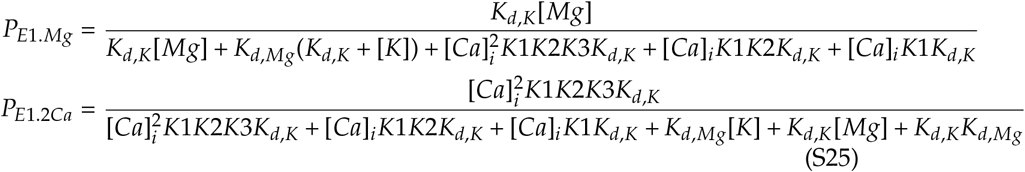

For Mg_−_K_−_, we have:

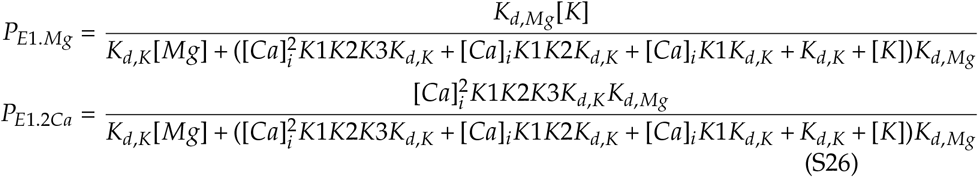

For Mg_+_K_+_, we have:

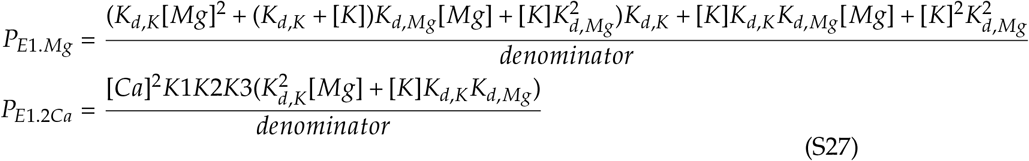

where denominator is:

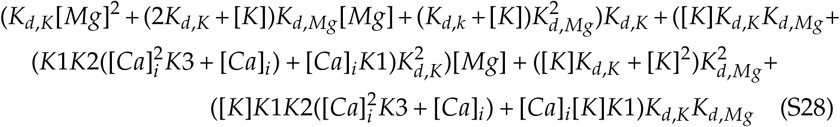

### S1.4 Supplementary Results

As shown in Table S5, the cations in the simulation maintained most of coordinating amino acids found in the crystal structure with few exceptions. In the crystal structure, site I Ca^2+^ has five coordinating amino acid oxygens within 2.5 Å. In our simulations, site I Ca^2+^ lost coordination with the N768 O_*δ*1_ atom (4.82 Å) and the coordination with T799 O_*γ*1_ also presented larger distances (3.46 ± 0.92 Å). However, the site I Ca^2+^ also gained two new coordinating amino acid oxygens with E771 O_*δ*2_ and A305 O at 2.20 and 2.27 Å, respectively. The net effect is that site I Ca^2+^ in simulation has slightly larger number of amino acid oxygen than WT (5.5 vs 5), as indicated in Fig. 4. Interestingly, in simulations site I Ca^2+^ also gained one more coordinating water when compared with crystal structure, making the number of coordinating waters reaching three. These waters were stabilized via hydrogen bonds with neighbouring oxygens, e.g., backbone oxygen of E908, N768 and the side chain oxygen of T799.

**Table S5:**
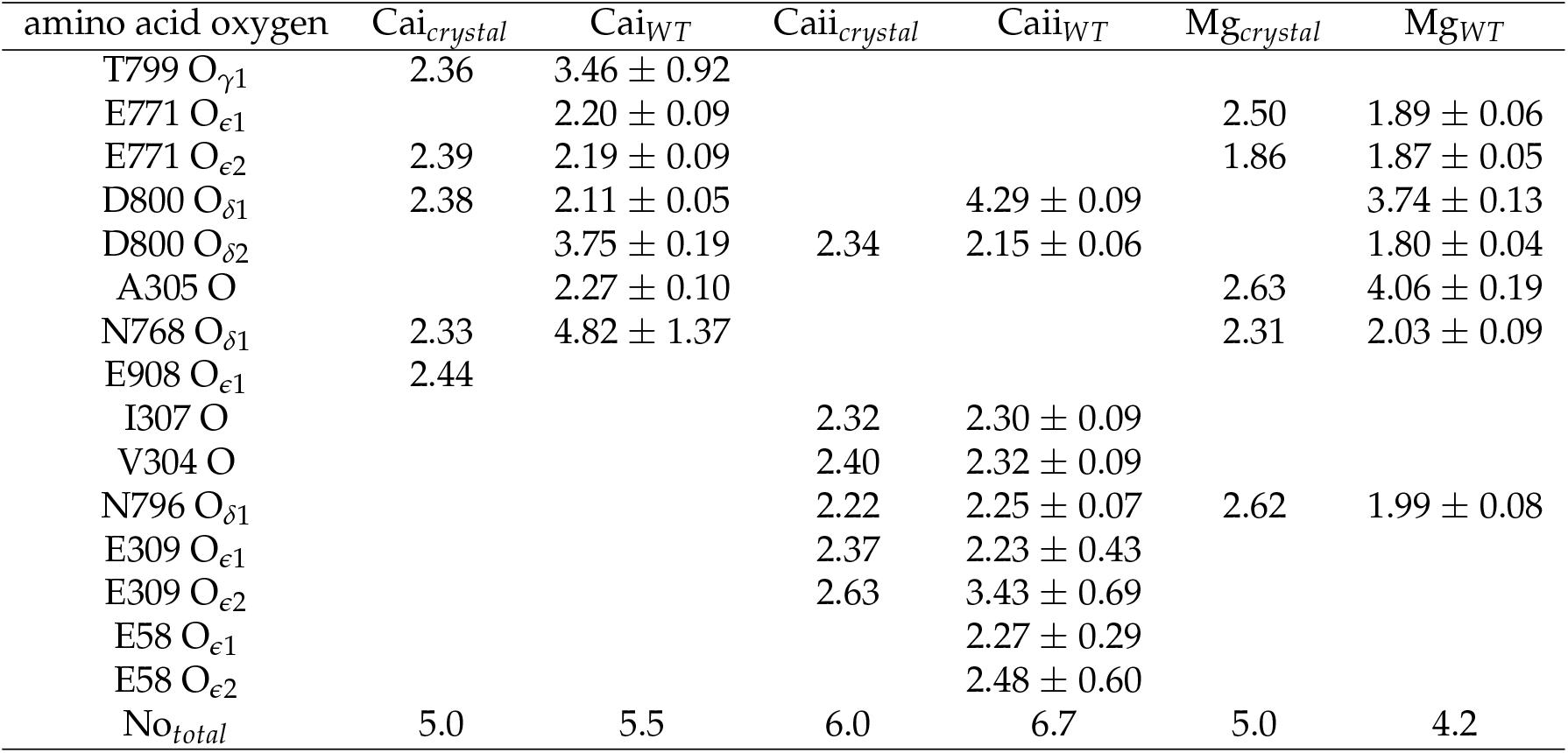
Distance between cation and coordinating amino acid oxygens in Å.

Meanwhile, the site II Ca^2+^ lost one coordination pairing with E309 O_*ϵ*2_ as the distance became 3.43 Å compared with 2.63 Å in crystal structure, whereas the ion established a bidentate coordination with E58. We note that the standard deviation between E58 O_*ϵ*2_ atom and Ca^2+^ is 0.60 Å, which is about six fold higher than the other positions, which indicated that E58 coordination is highly dynamic in comparison. For the E309Q and N796A variants, the number of coordinating amino acid oxygens to cations are comparable with the WT case but coordinating water density are significantly smaller (see inserted bar graphs in Fig. 4). Site II Ca^2+^ preserved coordination reflected in crystal structure. Similarly, Mg^2+^ lost coordination with A305 O but gain coordination with D800 O_*δ*2_ and therefore maintained same coordination number as crystal structure.

At the time of this study, to our knowledge there is no experimentally-determined structural information of SERCA with K^+^ bound at transmembrane sites, however, it has been reported that K^+^ binds to site I with submillimolar affinity [18]. In this study we show that the initially K^+^ placed at binding site II is unstable and exchange frequently with K^+^ ions in solvent at a sub-100 ns timescale while K^+^ at site I is stable as shown in Fig. S6. In the simulations, K^+^ at site I forms three stable coordination pairs with E771 O_*ϵ*1_, O_*ϵ*2_ atoms and T799 O_*γ*1_ atom at around 3.0 Å and a highly dynamic coordination with D800 O_*δ*2_ atom (3.99 ± 1.80 Å) as indicated by the large standard deviation, therefore yielding a total number of coordinating amino acid oxygens of approximately 3.9. We conclude that the MD simulations concur with the experimentally-resolved coordination patterns, although the identity of the coordinating residues present an appreciable degree of variability. We later relate these coordination pattern to binding potential.

In order to achieve more thorough understanding regarding energetic contribution from water, we analyzed thermodynamic properties of coordinating water sites during MD via GIST model. As shown in Fig. S14, around site I Ca^2+^, WT has three groups of water sites with −12 kcal/mol stability while N796A has one and E309Q has almost zero. These three groups of water sites in WT satisfy coordination distance with Ca^2+^ and correspond to the three coordinating water molecules as shown in Fig. 4, implying that these water coordinations considerably enhance Ca^2+^ affinity at site I in WT case. We also reported the GIST results for site II Ca^2+^ as shown in Fig. S8. In WT and E309Q cases, these is no stable water sites within appropriate distance to Ca^2+^, however, for N796A, one group of water sites with −12 kcal/mol stability presents and stabilizes Ca^2+^ at around 3.3 Å. Since the numbers of amino acid oxygens for WT/E309Q are comparable and greater than N796A, the additional stable water site group in N796A could make the site II Ca^2+^ binding affinity comparable among WT and variants. Therefore, it is evident that both coordinated amino acid oxygens and bound waters contribute to cation affinities in WT and mutant SERCA. Further, coordinated waters may compensate for the inability to optimally bind cations in sites I and II, owing to structural limitations.

**Figure S14:**
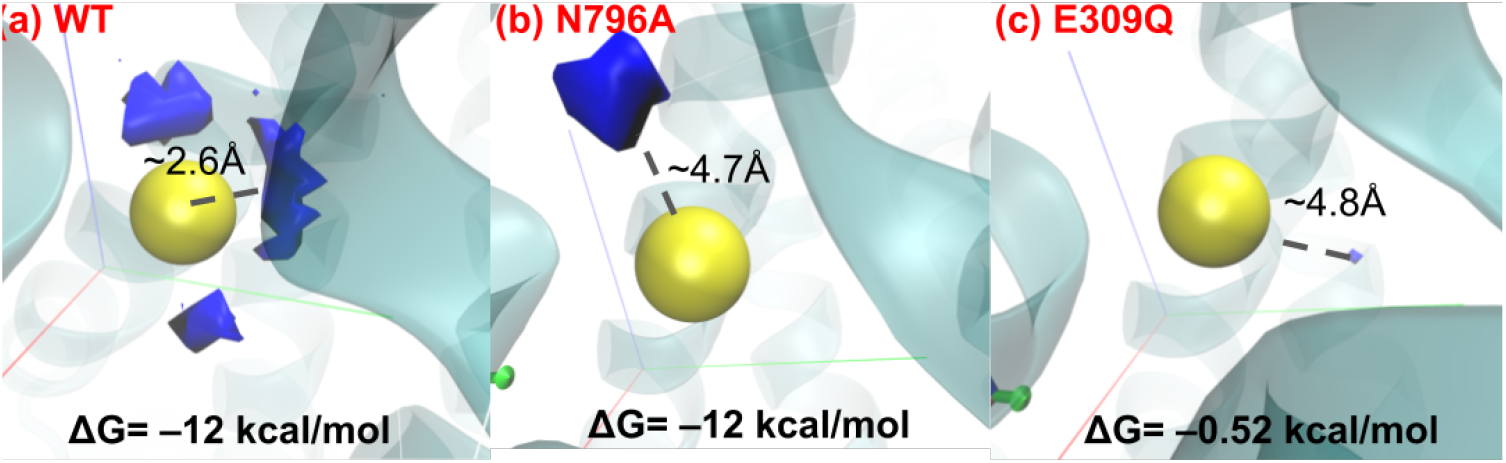
Free energy of water sites around site I Ca^2+^ calculated by GIST. For WT and N796A, the blue isosurface corresponds to water sites with Δ*G* = −12 kcal/mol, while for E309Q the middle value of Δ*G* (range is −1.03 to 0 kcal/mol) is shown. The distance between Ca^2+^ and isosurface is also shown. The site II Ca^2+^ data is in Fig. S8.

